# Skin and Gut Microbiome Features Associated with Resistance to *Corynebacterium bovis*-Associated Disease in Nude Mice (*Mus musculus*)

**DOI:** 10.64898/2026.09.15.751397

**Authors:** Kate E. Fodor, Amanda C. Ritter, Kourtney Nickerson, Aiswarya Prasad, Rodolfo J. Ricart Arbona, Janina Krumbeck, Neil S. Lipman

## Abstract

*Corynebacterium bovis* is an important opportunistic pathogen of immunodeficient mice and the etiologic agent of *Corynebacterium*-associated hyperkeratosis (CAH). Although disease severity varies, specific vendor-derived microbiomes have been shown to protect against CAH. Preinoculation with nonpathogenic *Corynebacterium amycolatum* prior to *C. bovis* infection has been demonstrated to limit disease severity. To define community features associated with protection, axenic outbred athymic nude mice stocks (A, B, and C) were reassociated with donor microbiomes from four vendor sources (A1, A2, B, and C) and topically challenged with *C. bovis*. *C. amycolatum* was added to the A1 microbiome in a separate group. Skin (Stock A) and fecal (Stocks A, B, and C) microbiomes were evaluated at 21 days post-inoculation via shotgun metagenomic sequencing. The A2 microbiome, previously associated with resistance to clinical disease and minimal skin pathology, exhibited greater cutaneous microbial evenness after challenge and significantly lower relative abundance of *C. bovis* than all other microbiome groups. Addition of *C. amycolatum* to the A1 microbiome did not fully confer disease protection and relative abundance was low, suggesting that sustained colonization by this organism was insufficient to explain the protective phenotype seen with the A2 microbiome. *C. kroppenstedtii* was detected in all A2 skin samples and was absent from all other groups. Several anaerobic taxa, including *Duncaniella dubosii, D. muris, Bacteroides caecimuris*, and *Muribaculum gordoncarteri*, were uniquely detected or enriched in the A2-associated microbiome, whereas *Mammaliicoccus lentus* and *Staphylococcus nepalensis* were absent from A2 but present in all nonprotective groups. Gut alpha diversity did not differ significantly among microbiomes, although several taxa, including segmented filamentous bacteria, were enriched in A2 feces. These findings associate resistance to CAH with preservation of cutaneous community structure, suppression of *C. bovis* prominence, and distinct microbial taxa. The identified organisms represent candidates for future mechanistic studies of microbiome-mediated colonization resistance.

## Introduction

*Corynebacterium bovis* is a highly contagious, gram-positive opportunistic skin pathogen of athymic nude and other immunodeficient mice and is the etiologic agent of Corynebacterium-associated hyperkeratosis (CAH), also known as scaly skin disease.^1–3^ Disease expression is variable and depends on factors including mouse age, bacterial isolate, and inoculum dose.^1,4^ In susceptible mice, infection may cause erythema, adherent keratinaceous scale, hyperkeratotic dermatitis, alopecia or rough hair coat in hirsute immunodeficient strains, weight loss or reduced body condition, dehydration, increased grooming or pruritus, lethargy, and hunched posture; histologic lesions typically include orthokeratotic hyperkeratosis, acanthosis, inflammation, and intracorneal gram-positive bacterial colonies.^1,4–6^ Once introduced into a facility, *C. bovis* is difficult to contain or eradicate because the organism can persist in the environment and transmission may occur through contaminated bedding, keratin flakes, fomites, husbandry procedures, aerosols generated during cage change, and biologic or research materials.^3,7^ Recent institutional surveys indicate that *C. bovis* remains common in U.S. academic and cancer-center vivaria despite increased surveillance and bioexclusion efforts.^8^ Antibiotic-based management has had context-dependent success. Prophylactic or metaphylactic approaches may suppress infection or transmission under some conditions, but treatment has not reliably eliminated *C. bovis* from infected colonies, clinical signs may recur after withdrawal, and antibiotic exposure can confound experimental outcomes by perturbing host-associated microbiota.^9,10^ In NSG mice, amoxicillin treatment has also been associated with marked intestinal microbiota changes, *Clostridioides difficile* colonization, and death.^11^ As a result, institutions face the difficult decision of pursuing complete bioexclusion through depopulation and sanitation or managing *C. bovis* as an endemic pathogen with its associated disease.^8^

The microbiome of animals and humans has become an increasingly important focus of biomedical research. Advances in molecular sequencing have enabled more detailed characterization and monitoring of complex microbial communities. The microbiome is now recognized as an important contributor to multiple aspects of host health, including immune development, metabolism, and disease susceptibility.^12–14^ In rodents, experimental manipulation of the microbiome can alter disease phenotypes, model outcomes, and behavior.^15,16^ Given its potential to affect experimental reproducibility, microbiome status should be considered, and when possible controlled or reported, in experimental design and interpretation.^17–19^

In addition to the variables described above, CAH severity is influenced by host genetics and microbiome composition.^2,20,21^ In a recent study, three distinct stocks of axenic outbred athymic nude mice were reassociated by cohousing with microbiome donors from commercial vendors and subsequently exposed to a pathogenic isolate of *C. bovis*.^21^ Mice from all three stocks failed to develop typical CAH lesions when reassociated with one specific donor microbiome, suggesting that microbiome composition can modulate disease expression. However, that study did not define the microbial features distinguishing protective from nonprotective reassociated communities. Because the microbiome donors were obtained from different commercial vendors and/or vendor colonies, and vendor-associated differences in murine microbial communities are documented, further characterization of these communities is warranted.^22–24^ Relatively little is known about the skin and gut microbiomes of immunocompromised mice. When described, reported skin-associated commensal or pathobiont taxa include corynebacteria, staphylococci, and *Lachnospiraceae*, among others.^10,25,26^ Following *C. bovis* infection, this organism can become a dominant member of the skin microbiome, reaching up to 90% relative abundance in athymic nude mice.^4^ Together, these observations warrant further investigation of *C. bovis*-microbiome interactions at both cutaneous and gastrointestinal sites.

*Corynebacterium amycolatum* has emerged as a candidate commensal that may influence susceptibility to *C. bovis*-associated hyperkeratosis. In a study examining the effects of athymic nude mouse stock and colony source on CAH severity, colonies in which *C. amycolatum* was detected on the skin exhibited minimal or absent clinical disease following *C. bovis* challenge, whereas colonies lacking detectable *C. amycolatum* developed more pronounced clinical disease.^20^ These observations suggested that *C. amycolatum* or other components of the associated skin microbiome might contribute to resistance to CAH, although host genetic differences and other microbiome features could not be excluded. More recently, direct topical exposure to *C. amycolatum* was evaluated as a potential strategy for mitigating *C. bovis*-associated disease.^21^ Pretreatment with *C. amycolatum* delayed disease onset and reduced peak clinical scores following *C. bovis* challenge, but did not prevent histopathologic changes and did not reproduce the degree of protection observed with a nonpathogenic *C. bovis* isolate.^2,21^ These findings suggest that *C. amycolatum* may have context-dependent effects on *C. bovis* colonization or disease expression and raise the possibility that its association with protection reflects interactions with other members of the skin microbiome rather than an autonomous protective effect.

In the present study, we evaluated the skin and fecal microbiomes of axenic outbred athymic nude mice reassociated with microbiomes from three commercial vendors subsequently inoculated with *C. bovis*. Our objective was to identify microbial community features associated with protection from CAH and to determine whether select organisms could account for disease susceptibility. Because previous observations implicated *C. amycolatum* as a potential protective skin commensal, we additionally evaluated the effect of topical *C. amycolatum* exposure within a nonprotective microbiome. We hypothesized that mice with reduced clinical disease would have lower relative abundance of *C. bovis* and that protective microbiomes would contain, or lack, specific microbial taxa associated with resistance to CAH. These findings may improve our understanding of *C. bovis*-microbiome-host interactions and help identify candidate organisms for future studies aimed at defining microbiome constituents that confer protection against CAH.

## Materials and Methods

### Experimental Design

Axenic F1 homozygous nude mice from three outbred athymic nude stocks: Stock A [Hsd:Athymic Nude-*Foxn1nu*], Stock B [J:NU], and Stock C [Crl:NU(NCr)-*Foxn1nu*] were used as experimental animals. At 4 to 6 weeks of age, male and female mice from each stock were randomly assigned to microbiome reassociation groups (n=8) and were cohoused for 14 days with conventionally raised athymic nude microbiome donor mice purchased directly from one of four commercial vendors and/or sites: A1, A2, B, or C. Sites A1 and A2 represented geographically distinct colonies from the same vendor.

The experimental design included 12 fully crossed stock-microbiome arms, corresponding to 3 mouse stocks x 4 donor microbiomes. In addition to directly evaluate the effect of *C. amycolatum*, an additional cohort (n=8) was cohoused with site A1 donor mice where all mice were topically inoculated with 10 CFU *C. amycolatum* isolate 25-1107-1 at the start of cohousing. *C. amycolatum* is not a constituent of the A1 microbiome. The *C. amycolatum*-augmented A1 arm was performed only in Stock A mice, yielding 13 stock-by-microbiome experimental arms in total.

After the 14-day cohousing period, donor mice were removed. Baseline aerobic skin cultures were collected from recipient mice for *C. bovis* and, as necessary, *C. amycolatum*. Fecal samples were also collected from each recipient mouse prior to *C. bovis* inoculation. Recipient mice, now 6 to 8 weeks old, were then topically inoculated with either 10 CFU of pathogenic *C. bovis* isolate 7894 (n=6/group) or sterile culture medium (n=2/group) as a negative control. At the experimental endpoint 21 days post-inoculation (dpi), an additional aerobic skin culture, skin swab, and fresh fecal pellets were collected from each mouse. Sex was balanced within each arm, with 3 male and 3 female mice in the *C. bovis*-challenged group and 1 male and 1 female mouse in the media-control group. Across the 13 experimental arms, this yielded 78 *C. bovis*-challenged mice and 26 media-control mice, for a total experimental sample size of 104 mice. Clinical data and skin histopathology were presented previously.^21^

### Animals

Heterozygous female (*Foxn1nu/+*) and homozygous male (*Foxn1nu/nu*) outbred athymic nude mice, 5 to 8 weeks of age, were purchased from three commercial vendors to generate axenic mouse stocks: Crl:NU(NCr)-*Foxn1nu* from Charles River Laboratories, Wilmington, MA; Hsd:Athymic Nude-*Foxn1nu* from Inotiv, Livermore, CA; and J:NU from The Jackson Laboratory, Bar Harbor, ME. Each stock was rederived to the germfree state by cesarean rederivation. Axenic F0 breeding pairs were then established for each stock to generate F1 experimental animals. The germfree status of each litter was confirmed at weaning by 16S PCR (IDEXX, Westbrook, ME; fecal sample).

Homozygous, axenic nude F1 mice were identified by phenotype and subsequently inoculated with *C. bovis* or sterile media at 6 to 8 weeks of age. Equal numbers of male and female mice were used across experimental groups. The total experimental sample size was 104 mice.

Conventionally raised athymic nude mice were purchased for use as microbiome donors from sites A1 (Livermore, CA), A2 (Denver, PA), B, and C and maintained in sterile (caging, food and water), hermetically sealed, individually ventilated, HEPA-filtered, isolator cages upon receipt to preserve microbiome integrity. Donor mice were screened for *C. bovis* and *C. amycolatum* by aerobic culture before cohousing; site A1 donors lacked detectable *C. amycolatum*, whereas site A2 and B donors were colonized with *C. amycolatum* as part of their endogenous microbiomes. All donor mice were *C. bovis* free.

All mice were free of Sendai virus, pneumonia virus of mice, mouse hepatitis virus, minute virus of mice, mouse parvovirus, Theiler meningoencephalitis virus, reovirus type 3, epizootic diarrhea of infant mice (mouse rotavirus), murine adenovirus, polyoma virus, K virus, murine cytomegalovirus, mouse thymic virus, lymphocytic choriomeningitis virus, Hantaan virus, ectromelia virus, lactate dehydrogenase elevating virus, *Bordetella bronchiseptica*, *Citrobacter rodentium*, *Clostridium piliforme, Corynebacterium bovis, Corynebacterium kutscheri, Filobacterium rodentium, Mycoplasma pulmonis, Salmonella* spp*., Streptobacillus moniliformis, Streptococcus pneumoniae, Encephalitozoon cuniculi,* ectoparasites, endoparasites, and enteric protozoa. Mice received directly from vendors and housed in isocages were also free of mouse norovirus, rodent chaphamaparvovirus (except Inotiv), mouse thymic virus, *Helicobacter* spp, *Klebsiella pneumoniae*, *Klebsiella oxytoca*, *Pasteurella multocida*, *Rodentibacter pneumotropicus*, *Rodentibacter heylii*, *Pneumocystis murina*, *Proteus mirabilis*, *Pseudomonas aeruginosa*, *Staphylococcus aureus*, and beta-hemolytic *Streptococcus* spp. Based on health monitoring reports for the Inotiv PA site, mice from this location were not confirmed free of murine adenovirus, polyoma virus, Hantaan virus, ectromelia virus, lactate dehydrogenase elevating virus, mouse kidney parvovirus, mouse thymic virus, *B. bronchiseptica*, *F. rodentium*, *S. moniliformis*, *E. cuniculi*, and *P. multocida*.

### Husbandry and housing

Prior to rederivation, mice were housed in individually ventilated, autoclaved, polysulfone cages with stainless steel wire bar lids and filter tops (no. 9; Thoren Caging Systems, Hazleton, PA) on autoclaved aspen-chip bedding (PWI Industries, Quebec, Canada) with up to 3 mice per cage. Each cage was provided an enrichment bag made from Glatfelter paper containing 6 g of crinkled paper strips (EnviroPak; WF Fisher and Son, Branchburg, NJ). Each cage (including cage bottom, bedding, wire bar lid, and water bottle) was changed weekly within a vertical-flow cage changing station (AireGard NU-S619-400, NuAire, Plymouth, MN). Mice were fed a closed-source, natural ingredient, flash-autoclaved, irradiated diet (5053; LabDiet) and provided *ad libitum* reverse osmosis acidified (pH 2.5 to 2.8 with hydrochloric acid) water in polyphenylsulfone bottles with autoclaved stainless-steel caps and sipper tubes (Tecniplast, West Chester, PA).^27^

Following rederivation, mice were housed in autoclaved, solid bottom and top, gasketed and sealed, polysulfone, positive-pressure, individually ventilated isolator cages (Sentry SPP Mouse; Allentown Caging Equipment, Allentown, NJ) with autoclaved aspen chip bedding (PWI Industries) with up to 6 mice per cage. Cages were clearly labeled with stock, vendor site, sex, and experimental group. HEPA-filtered air (filtration at rack and cage level) ventilated each cage at approximately 30 air changes per hour. The HEPA-filtered rack effluent vented directly into the building’s HVAC system. Mice received γ-irradiated, autoclaved feed (5KA1, LabDiet) and non-acidified, autoclaved reverse osmosis water *ad libitum*. Each cage was provided with a sterile bag of Glatfelter paper containing 6 g of crinkled paper strips (EnviroPAK^â^, WF Fisher and Son) for enrichment. Microbiome donors and recipients were housed in isocages as described for axenic mice to maintain their specific microbiome. Cages housing axenic mice were changed at least every 8 weeks and every 2 weeks for all others. All manipulations, including cage changes, were performed in a horizontal laminar flow hood (AireGard ES NU-340, NuAire, Plymouth, MN) using sterile gloves and aseptic technique. To minimize cross-contamination, germfree and control mice were handled before other animals, with *C. amycolatum*- and *C. bovis*-inoculated mice handled last. The animal room was ventilated with 100% fresh air at a minimum of 10 air changes hourly and maintained at 72 ± 2°F (21.5 ± 1°C), relative humidity between 30% and 70%, and a 12:12 h light-dark cycle (lights on at 0600, off at 1800). All animal use was approved by Memorial Sloan Kettering’s (MSK) IACUC and conducted in accordance with AALAS’s position statements on the Humane Care and Use of Laboratory Animals and Alleviating Pain and Distress in Laboratory Animals. MSK’s animal care and use program is AAALAC-accredited and operates in accordance with the recommendations provided in the *Guide for the Care and Use of Laboratory Animals* (8th ed.).^28^

### Caesarian rederivation

Heterozygous athymic nude female mice from each stock were bred to a homozygous male nude mouse, examined for copulatory plugs, and weighed weekly to identify pregnancies. Timed-pregnant animals received 4.5 mg SQ medroxyprogesterone acetate (Mylan, Canonsburg, PA) on gestational day 17.5 to inhibit parturition. On gestational day 19.5, an aseptic field was prepared in a horizontal laminar flow hood (AireGard ES NU-340, NuAire, Plymouth, MN). Each pregnant mouse was euthanized by cervical dislocation and fully submerged for 1 minute in 0.5% hydrogen peroxide (RTU Peroxigard, Virox Technologies Inc., Ontario, CA) before placement on a sterile field for a hysterectomy. The abdomen was incised, the uterus was removed *en bloc* and submerged in chlorine dioxide disinfectant solution (Clidox [1:4:1], Pharmacal Laboratories, Naugatuck, CT) for 30 seconds. The uterus was placed on a sterile heated field, each uterine horn was incised, the pups were removed and stimulated with sterile gauze until spontaneous breathing occurred. All healthy pups were transferred aseptically to an isocage containing an axenic BALB/c foster dam and 2 of the foster dam’s pups whose age was within 4 days of the litter being fostered. The pups’ germfree status was confirmed at weaning via negative 16S PCR of fecal pellets.

### Establishment of unique stock/microbiome combinations by cohousing

Conventionally reared female athymic nude mice imported directly from sites A1, A2, B, and C served as microbiome donors. Microbiome donors were confirmed negative for *C. bovis* and, for site A1, *C. amycolatum* via aerobic culture. Only female mice were used as microbiome donors to minimize intracage mouse aggression.

Axenic mice of each stock (up to 4 per cage) were each cohoused for 2 weeks with a microbiome donor mouse from each of the 4 vendor sites, allowing sufficient time for microbiome reassociation through coprophagy and grooming. One experimental group of mice that was cohoused with an A1 microbiome donor also received 10 CFU of *C. amycolatum* 25- 1107-1 topically at the time of cohousing. On arrival and between cohousing periods, microbiome donor mice were housed in isolator cages and handled aseptically to maintain their unique microbiome.

### Corynebacterium bovis and Corynebacterium amycolatum propagation and inoculation

*C. bovis* isolate 7894 and *C. amycolatum* isolate 25-1107-1 stored as frozen stocks were grown on trypticase soy agar supplemented with 5% sheep blood (BBL TSA II 5% SB; Becton Dickinson, Sparks, MD) at 37°C with 5% CO□ for 48 h. Growth curves for the isolates were previously established.^1,2^ *C. bovis* isolate 7894 originated at MSK and was shown to be the most pathogenic of 6 *C. bovis* isolates evaluated in a study evaluating 9 *C. bovis* isolates.^1,5^ *C. amycolatum* 25-1107-1 originated from a *C. bovis*-negative mouse at MSK with clinical signs similar to CAH. As the infectious dose for this isolate was not established, a high inoculation dose was used to ensure colonization. Inocula were prepared from bacterial suspensions during the midlog growth phase, titrated and serially diluted in brain heart infusion broth (BHI; Becton Dickinson) supplemented with 0.1% Tween 80 (VWR Chemicals, Solon, OH) to obtain 10 CFU (*C. bovis*) and 10□ CFU (*C. amycolatum*) ± 15% bacteria in 50 μL of media. Bacterial concentrations were estimated with a MacFarland densitometer.^1^ Sterile 2-mL polypropylene tubes (Fisher Scientific, Waltham, MA) containing the inocula were transferred to the vivarium on ice, and animals were topically inoculated within 2 hours. All animal manipulations were performed in a horizontal laminar flow hood (AireGard ES NU-340, NuAire, Plymouth, MN) using sterile materials and aseptic technique. After inoculation, the concentration of each inoculum was confirmed and enumerated by plate count.

Control mice, inoculated with sterile media, were handled before animals inoculated with *C. bovis*. Each cage was removed from its rack, sprayed on all sides with disinfectant (Peroxigard [1:16]; Virox Technologies, Ontario, CA) and placed in the laminar flow hood. Each cage was opened for less than 5 min. Each mouse was gently grasped at the base of the tail, slightly elevating its hindquarters, and the bacterial inoculum was applied directly to the dorsal midline using a sterile filter micropipette (P200N; Marshall Scientific, Hampton, NH). A 50-μL bacterial suspension or bacterial-free BHI broth (negative controls) was deposited at 4 to 5 sites along the skin using a sterile filtered 200-μL pipette tip (Filtered Pipet Tips; Crystalgen, Commack, NY), delivering 10 to 15 μL per site to prevent immediate runoff. Cages were reassembled and returned to the rack, and new sterile gloves were donned between each cage.

### Skin sample collection and microbiome sequencing

Skin samples for microbiome sequencing were aseptically collected from only Stock A mice at the experimental endpoint by rubbing an individually packaged adhesive swab (PurFlock ULTRA, Puritan Medical Products Company LLC, Guilford, ME) 5 times along the left flank from the base of the tail cranially to the point of the shoulder and caudally back to the tail base. Swab tips were broken off into sterile 2-mL polypropylene centrifuge tubes and stored at -80°C within 4 hours until analyzed. Swabs were shipped on dry ice to the testing facility (MiDOG LLC, Tustin, CA). The negative control samples for this stock were included to provide information regarding the microbiome composition without the influence of *C. bovis*.

Genomic DNA was extracted using a microbial DNA extraction kit (ZymoBIOMICS^®^ DNA Microprep Kit D4301, Zymo Research, Irvine, CA). The kit was selected for use with skin swab samples due to its low elution volume yielding more concentrated DNA. Sequencing library preparation and data analysis for shotgun metagenomic sequencing were performed using a DNA preparation kit (Illumina DNA Prep Kit, Illumina, San Diego, CA) with 10 bp unique dual indexes. Libraries were quantified (Qubit, ThermoFisher Scientific), pooled by equal abundance, and the final pool quantified by qPCR. Sequencing was performed by next generation sequence analyzers (Illumina NextSeq^®^ 2000 or NovaSeq^®^ X).

Raw reads were trimmed with Trimmomatic-0.33 using a 6 bp sliding window with quality cutoff of 20; reads shorter than 70 bp were discarded. Host-derived reads were removed with Kraken2 against common eukaryote host genomes including *Mus musculus*. Low-complexity reads were filtered with sdust.^29,30^

Microbial composition was profiled using Sourmash with abundance-aware DNA sketches generated at k=51 and scaled=1000. Metagenomic classification was performed with sourmash gather using the parameters ksize 51 and threshold-bp 5000 against GTDB species representative database RS207 for bacteria and pre-formatted GenBank databases (v. 2022.03) for viruses, protozoa, and fungi.^31^ At this sketch scale, the gather threshold corresponds to approximately five unique hashes, or ∼5 kb of estimated unique sequence evidence. Abundance was determined as the sum of abundance-weighted hashes found in the respective taxon and relative abundance determined as the fraction of query hashes uniquely matched (weighted by multiplicity) that are uniquely matched to taxon. Gather hits were subsequently filtered using the following parameters chosen in a data-dependent manner so as to retain meaningful amount of data: intersect_bp ≥ 10,000 bp (retains ∼93%), f_unique_to_query ≥ 1×10 (retains ∼97%), and n_unique_weighted_found ≥ 10 (retains ∼85%). Species-level count tables were assembled into a phyloseq object in R for downstream analysis. Eukaryotic and bacterial fractions were separated, only the latter was included in downstream analysis.

### Fecal sample collection and microbiome sequencing

Animals were grasped gently by the base of the tail and lifted slightly, allowing all 4 limbs to grasp the wire bar lid. Two fecal pellets were aseptically collected per animal per timepoint and placed directly into separate barcoded tubes containing DNA stabilization buffer (Transnetyx Microbiome; Transnetyx Inc., Cordova, TN). Fecal pellets were collected at approximately the same time of day to minimize variability (between 0800 and 1100). Samples were housed at room temperature until analysis.

Study endpoint fecal samples were assayed to assess changes in microbial populations across stock and microbiome. Male and female samples per cohort were aseptically pooled by sex to yield 2 samples per stock-microbiome combination, allowing microbiome comparison across all 3 stocks. The individual samples were vortexed to suspend the feces homogenously in the buffer. Using a sterile pipette, approximately a third of each homogenized sample was transferred to a new, empty microbiome tube (Transnetyx, Inc.). The 3 male samples and the 3 female samples for each microbiome-stock group were pooled together to create 1 combined male and 1 combined female sample for each microbiome-stock group. These two pooled samples per group were analyzed for fecal microbiome composition across all mouse stocks and microbiomes, to evaluate and identify major disparities and trends between the stocks and microbiomes. To simplify select comparisons, the “protective” microbiome refers to the A2 microbiome, due to the lack of clinical disease following inoculation with a pathogenic *C. bovis* isolate, and “nonprotective” refers to all other microbiome groups (A1, A1+Ca, B, and C).

Samples were shipped to a commercial laboratory for DNA extraction, library preparation, and sequencing (Transnetyx Inc.).^32^ DNA extraction was performed (Qiagen DNeasy 96 PowerSoil Pro QIAcube HT extraction kit, Quiagen Inc., Germantown, MD) and a protocol for reproducible extraction of inhibitor-free, high molecular weight genomic DNA that captures the microbial diversity of stool samples followed. Genomic DNA was converted into sequencing libraries using the KAPA HyperPlus library preparation protocol optimized for minimal bias. Unique dual-indexed adapters were used to ensure that reads and/or organisms were not misassigned. The libraries were sequenced (NextSeq 2000 instrument, Illumina, San Diego, CA) via the shotgun sequencing method at a depth of 2 million 2 × 150-bp read pairs to enable species and strain level taxonomic resolution. Raw data (in the form of FASTQ files) were analyzed using the One Codex analysis software and database (One Codex, San Francisco, CA), which consists of approximately 148 K complete microbial genomes, including 71 K distinct bacterial genomes, 72 K viral genomes, and thousands of archaeal, eukaryotic genomes, and mouse gut metagenome-assembled genomes. Human and mouse genomes are included to screen out host reads. The database was assembled from both public and private sources, with a combination of automated and manual curation steps to remove low-quality or mislabeled records. Every individual sequence (NGS read or contig) is compared against the One Codex database by exact alignment using k-mers where k = 31. Based on the relative frequency of unique k-mers in the sample, sequencing artifacts are filtered out of the sample to eliminate false positive results caused by contamination or sequencing artifacts. The relative abundance of each microbial species is estimated based on the depth and coverage of sequencing across every available reference genome.

### Bioinformatics and Statistical Analysis

For comparative analysis of skin microbiome data, observed species richness, Shannon entropy, and Simpson index (alpha diversity indices) were calculated from species-level count data (vegan and phyloseq packages) and pairwise group differences were assessed using Wilcoxon rank-sum tests (rstatix package). Pairwise sample dissimilarity (beta diversity) was computed using Jaccard distance (presence/absence) and Bray-Curtis dissimilarity (log-transformed counts + 1 pseudo-count) via the vegan package. Principal coordinates analysis (PCoA) was performed using phyloseq. Permutational multivariate ANOVA (PERMANOVA) was conducted using the adonis2 function from the vegan package (999 permutations) to test community-level effects by group. Adjusted R^2^ or ω values were calculated using the adonis_OmegaSq function from the MicEco package (https://github.com/Russel88/MicEco). A *P* value less than or equal to 0.05 was considered statistically significant. Relative abundance was computed per sample at the phylum, family, genus, and species levels. All relative abundances per sample were summed at the respective taxonomic level and normalized to 100% to compare across samples. Stacked bar plots were generated with ggplot2 and ggh4x, with samples grouped by microbiome source and stratified by infection status or as mentioned in the respective figure. Taxa outside the top-ranked groups were collapsed into an “Other” category.

Fecal microbiome analyses were conducted in R using a phyloseq object containing count data, taxonomic annotations, and sample metadata (phyloseq). Post-challenge samples were selected, and taxa with zero or very low counts were removed where appropriate (phyloseq). Alpha diversity was calculated from non-normalized counts using Observed richness, Shannon entropy, and Simpson index, then displayed as box plots with individual-animal points and separate metric panels (phyloseq, ggplot2); selected phenotype comparisons were evaluated using Wilcoxon rank-sum tests and displayed with significance brackets (ggsignif, R stats). Beta diversity was assessed using Jaccard distances and Bray-Curtis distances, followed by principal coordinate analysis (phyloseq); group differences were tested by PERMANOVA with 999 permutations (vegan) and visualized with 95% confidence ellipses (ggplot2). Relative abundance was calculated within each sample, aggregated at the phylum, genus, and species levels, and plotted as stacked bars representing individual animals (phyloseq, dplyr, ggplot2, scales); missing taxonomic assignments were retained as “Unclassified,” low-abundance taxa were collapsed into “Other,” and sample totals were verified to sum to one after aggregation. Corynebacteria were analyzed separately at the species level using both raw read counts and within-genus relative abundance (phyloseq, dplyr, ggplot2). Differential abundance was assessed from raw counts using negative-binomial generalized linear models (DESeq2), with low-count taxa removed and Benjamini-Hochberg-adjusted *P*-values ≤ 0.05 were considered significant. As microbiome source and protection status were partially confounded, valid protection comparisons were conducted within microbiome groups containing at least two outcome categories. Data reshaping, ordering, joining, and summary calculations used dplyr, tibble, and tidyr, while multi-panel figures were assembled using patchwork.

## Results

### Skin microbiome

At the experimental endpoint, skin samples from six *C. bovis*-infected and two *C. bovis*-free Stock A mice across five microbiome variations were analyzed for microbial composition. Trimmed sequencing reads showed an average of ∼29% host reads (Supplementary Figure S1). The observed alpha diversity between skin microbiomes was not significantly different between groups. However, Shannon (*P* ≤ 0.01) and Simpson (*P* ≤ 0.01) indices, used to evaluate community evenness, were significantly higher in the protective A2 microbiome as compared to all other microbiome groups in *C. bovis*-challenged mice (Figure 1B). The observed alpha diversity for negative control samples demonstrates relatively equal species diversity between microbiome groups (Figure 1A). Across all microbiome groups, *C. bovis*-infected samples had significantly decreased alpha diversity as compared to their negative controls (*P* ≤ 0.05) (Figure 2).

**Figure 1.**
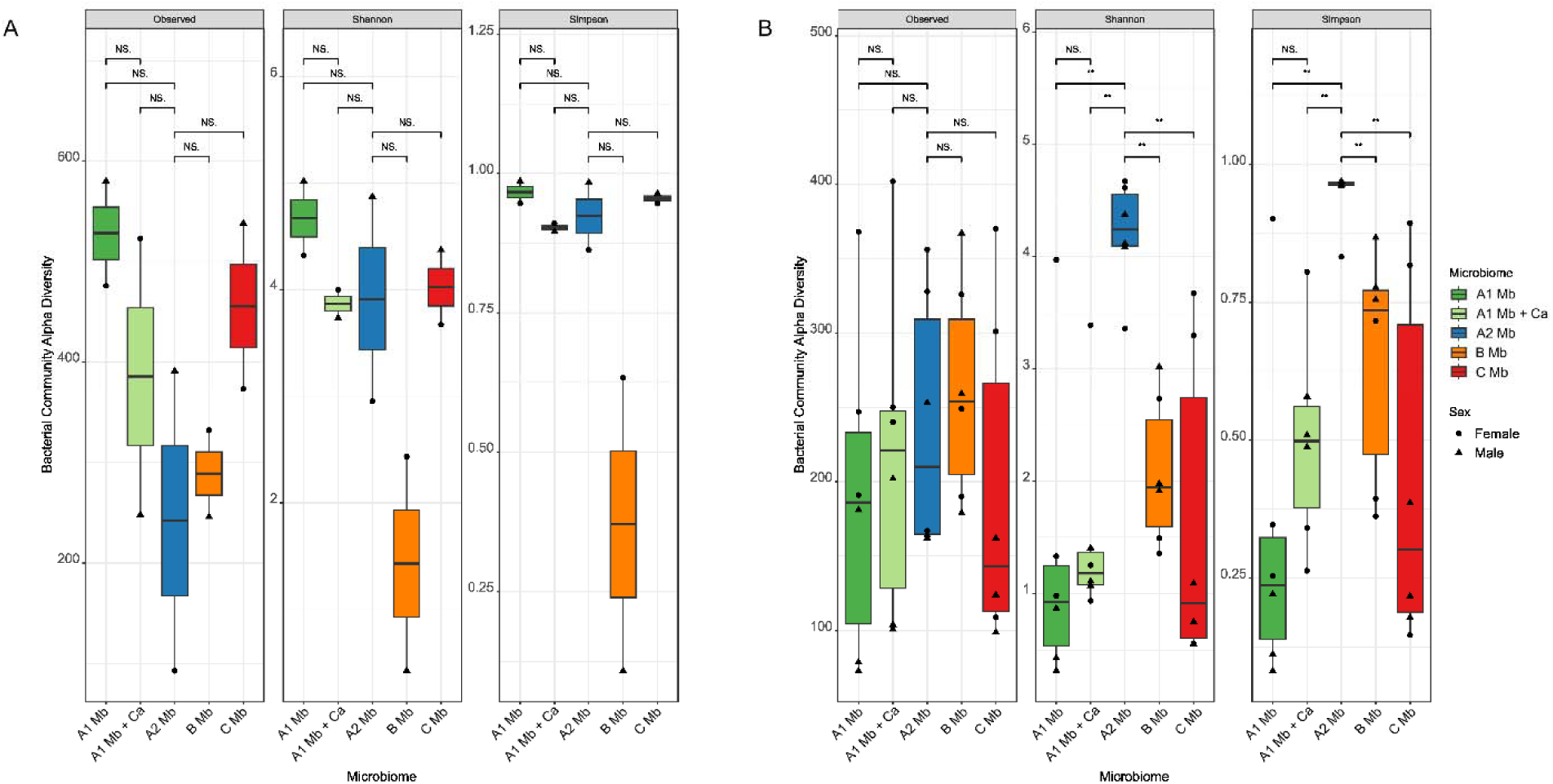
Comparison of cutaneous bacterial alpha diversity across microbiome groups and experimental conditions. Microbial community alpha diversity measured by Observed specie richness (left facet), Shannon diversity index (middle facet), and Simpson index (right facet) for Stock A mice across five microbiome profiles: A1 (dark green), A1 + *Corynebacterium amycolatum* (light green), A2 (blue), B (orange), and C (red). **(A)** Uninfected negative control baseline samples. **(B)** *Corynebacterium bovis*-infected samples at the experimental endpoint, showing a marked drop in diversity across non-protective groups and retention of higher community evenness in the protective A2 cohort. Individual data points represent individual mice, stratified by sex (female = circles, male = triangles). Box plots display the median (horizontal bar), interquartile range (IQR; box bounds), and minimum/maximum value (whiskers). NS, not significant; \*\**P* ≤ 0.01.

**Figure 2.**
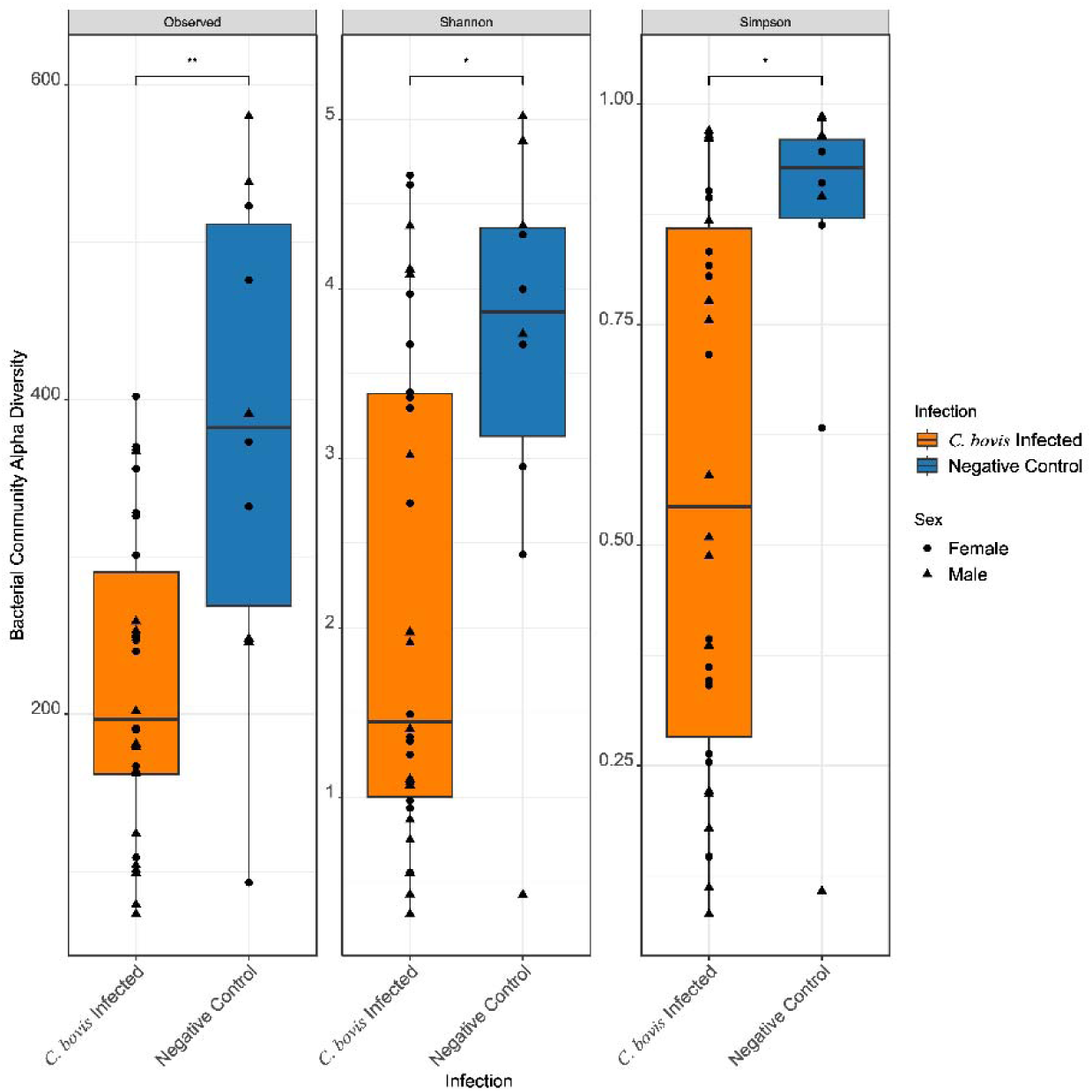
Reduction of cutaneous bacterial alpha diversity following *Corynebacterium bovis* infection. Comparison of skin microbial alpha diversity metrics: Observed species richness (left facet), Shannon diversity index (middle facet), and Simpson evenness index (right facet) between *C. bovis*-infected mice (orange) and uninfected negative controls (blue) across all microbiome groups at the experimental endpoint. Individual data points represent individual mice, stratified by biological sex (female = circles, male = triangles). Box plots depict the median (horizontal line), interquartile range (IQR; box limits), and minimum/maximum value (whiskers). There is significant overall depletion in bacterial species richness and community evenness in *C. bovis*-infected samples compared to uninfected controls. \**P* ≤ 0.05, \*\**P* ≤ 0.01.

Beta diversity metrics identified a significant relationship between the microbiome group and bacterial composition (ω² = 0.294 and 0.346). The PERMANOVA also detected significant microbiome community differences among the groups tested (*P*=0.001). In both the Jaccard and Bray-Curtis plots, the three Vendor A populations (A1, A1+Ca, and A2) showed distinct community overlap, indicating they are more closely related as compared to the microbiome of mice in the Vendor B and C groups (Figure 3). Interestingly, Vendor C’s negative control samples appeared to cluster within the 95% confidence interval of Vendor A2, where the *C. bovis*-infected samples from Vendor C were more disparate.

**Figure 3.**
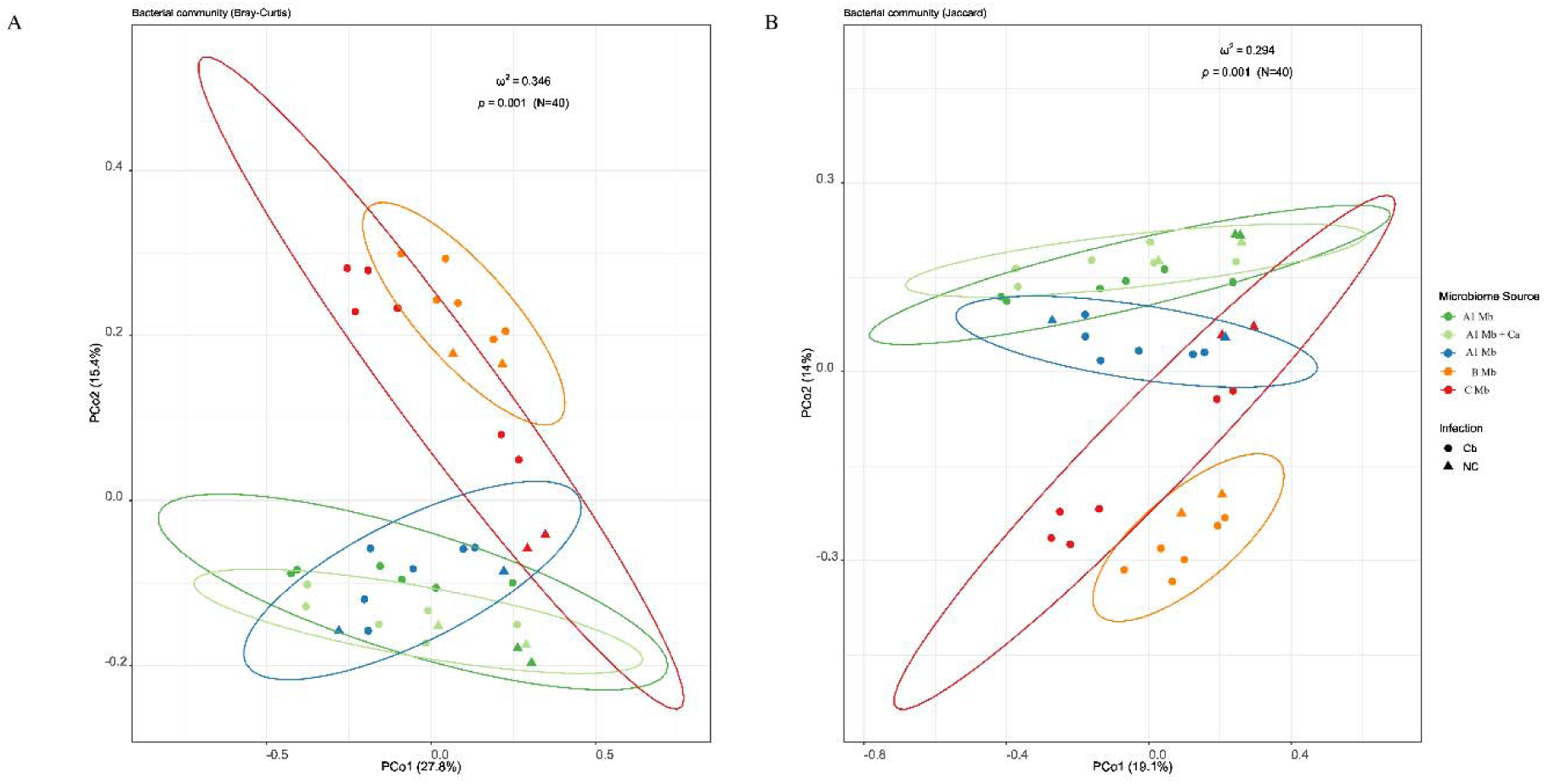
Cutaneous bacterial beta diversity across microbiome groups and infection status. Principal Coordinate Analysis (PCoA) plots of cutaneous bacterial communities based on **(A)** Jaccard distance (presence/absence) and **(B)** Bray-Curtis dissimilarity (relative abundance). Samples are colored by microbiome source: A1 (dark green), A1 + *Corynebacterium amycolatum* (light green), A2 (blue), B (orange), and C (red). Symbol shapes denote infection status: *Corynebacterium bovis*-infected (Cb; circles) and uninfected negative controls (NC; triangles). Ellipses represent 95% confidence intervals for each microbiome group. Community composition varied significantly by microbiome source across both metrics. PERMANOVA statistics ( and -values) are annotated within each panel (*P* = 0.01 for both metrics). There is marked spatial overlap among Vendor A populations (A1, A1 + *C. amycolatum*, and A2) compared to Vendors B and C, as well as the alignment of uninfected Vendor C samples near the A2 confidence interval in panel B.

Relative abundance profiles highlighted clear differences driven by microbiome source and infection status across multiple taxonomic levels (Supplementary Figures S2-S4). When restricting analysis to the genus *Corynebacterium*, composition and relative abundance varied substantially by microbiome (Figure 4). Negative control samples from groups A1 and C lacked endogenous *Corynebacterium* species entirely; upon infection, *C. bovis* accounted for 100% of the *Corynebacterium* reads in these groups. Conversely, the protective A2 microbiome harbored the greatest diversity of endogenous *Corynebacterium* species prior to infection. Although reduced following *C. bovis* challenge, this diversity persisted to a greater extent in A2 than in any other infected group (Figure 4). Groups A1+Ca and B contained fewer baseline *Corynebacterium* species than group A2. Furthermore, *C. amycolatum* was detected at low relative abundance in the A1+Ca group despite a high inoculation dose. Critically, the relative abundance of *C. bovis* post-challenge was significantly lower in A2 mice than in all other groups ( 0.01) and mean read count was markedly lower (Figure 5; Table 1).

**Figure 4.**
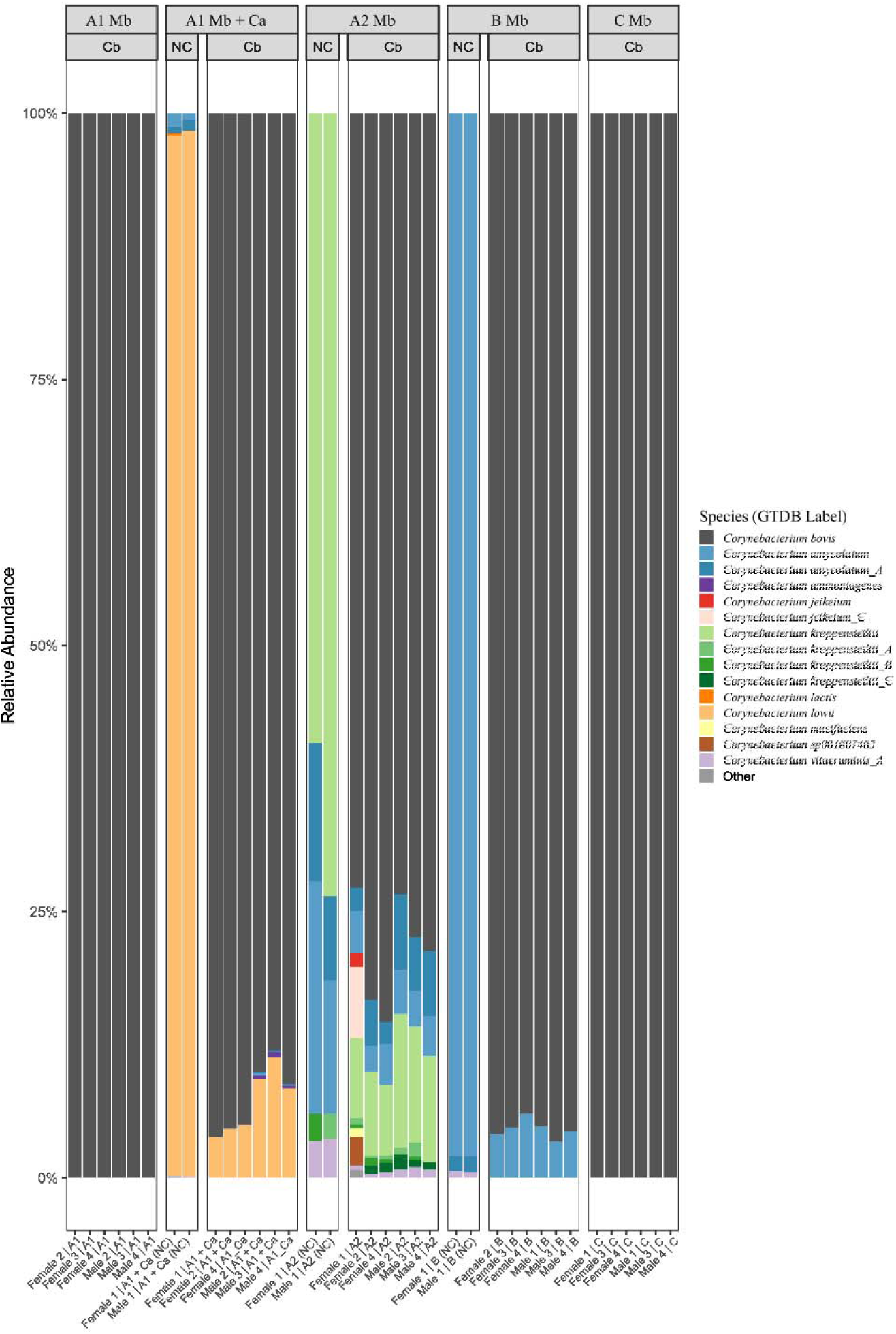
Relative abundance of cutaneous *Corynebacterium* species by microbiome group and infection status. Stacked bar plots depicting the taxonomic composition and relative abundance of *Corynebacterium* species on the skin across five microbiome profiles (A1, A1 + Ca [A1 + *C. amycolatum*], A2, B, and C) under uninfected negative control (NC) and *C. bovis*-infected (Cb) conditions. Each vertical bar represents a single animal sample. Individual *Corynebacterium* species and sub-species strains are identified by distinct colors, with *C. bovis* highlighted in dark gray. Microbiome groups without NC columns indicate that those groups lacked detectable *Corynebacterium* reads prior to *C. bovis* inoculation. Note the exclusive presence of endogenous *Corynebacterium* species including unique detection of *C. kroppenstedtii* lineages (greens/reds) in uninfected A2 samples, as well as the partial retention of non-*C. bovis* species diversity in A2 mice post-infection compared to near 100% *C. bovis* (dark gray) dominance in all other infected groups.

**Figure 5.**
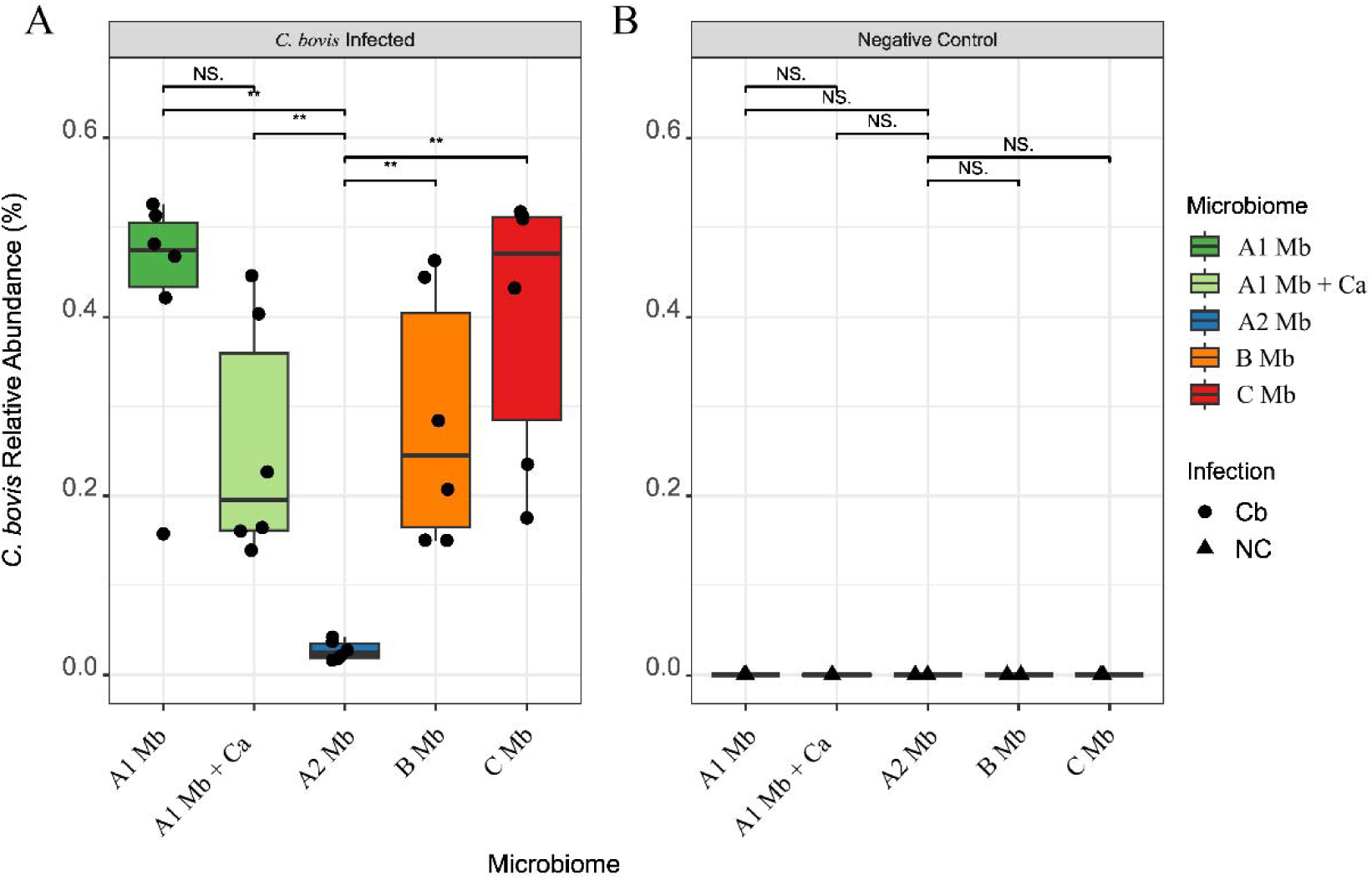
Significant suppression of *Corynebacterium bovis* relative abundance in the protective A2 microbiome. Relative abundance (%) of *C. bovis* on the skin of Stock A mice across five microbiome groups: A1 (dark green), A1 + Ca (A1 + *Corynebacterium amycolatum*; light green), A2 (blue), B (orange), and C (red). **(A)** *C. bovis*-infected samples at the experimental endpoint (circles). **(B)** Uninfected negative control baseline samples (triangles). Individual data points represent individual animals overlaid on box plots displaying the median (horizontal line), interquartile range (IQR; box limits), and minimum/maximum value (whiskers). The post-infection *C. bovis* relative abundance was significantly reduced in the A2 cohort compared to all other infected groups, while negative control samples exhibited no detectable *C. bovis* across all microbiome profiles. NS, not significant, \*\**P* ≤ 0.01.

**Table 1.** Mean raw sequencing read abundance (SD) of cutaneous *Corynebacterium bovis* by microbiome source.

| Microbiome Profile | Mean *C. bovis* Read Count ± SD |
| --- | --- |
| A1 | 88,790 ± 49,051 |
| A1+Ca | 104,629 ± 57,319 |
| A2 | 4,253 ± 2,375 |
| B | 152,351 ± 111,494 |
| C | 120,797 ± 52,365 |

To identify key members of the protective A2 skin flora, we screened for bacterial species uniquely present in this group. Species found in 100% of A2 samples but completely absent from all other groups were identified (Table 2). Uniquely, *Corynebacterium kroppenstedtii* was identified across all A2 samples, but was completely absent from all other groups (Figure 4); despite minor variability in strain-level taxonomic assignments, this species was consistently present in every A2 skin swab. Notably, this subset also included two anaerobic taxa typically associated with the mouse gut microbiome: *Duncaniella dubosii* and *D. muris*. Furthermore, several additional gut-associated anaerobes, including *Bacteroides caecimuris* and *Muribaculum gordoncarteri*, were present in ≥ 75% of A2 samples while remaining entirely absent from all other microbiome groups.

**Table 2.**
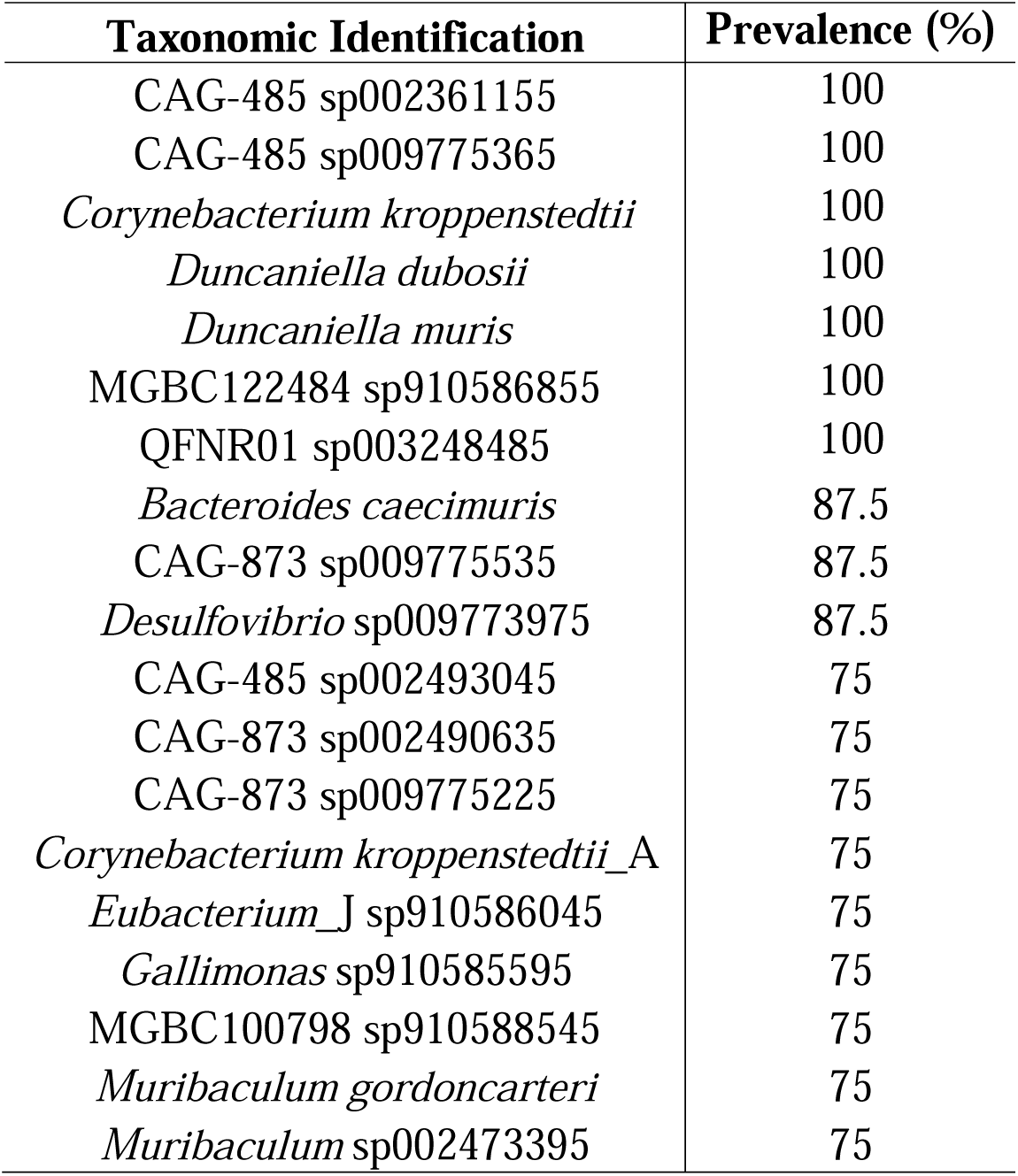
Cutaneous bacterial species uniquely detected in the protective A2 microbiome.

| **Taxonomic Identification** | **Prevalence (%)** |
| --- | --- |
| CAG-485 sp002361155 | 100 |
| CAG-485 sp009775365 | 100 |
| *Corynebacterium kroppenstedtii* | 100 |
| *Duncaniella dubosii* | 100 |
| *Duncaniella muris* | 100 |
| MGBC122484 sp910586855 | 100 |
| QFNR01 sp003248485 | 100 |
| *Bacteroides caecimuris* | 87.5 |
| CAG-873 sp009775535 | 87.5 |
| *Desulfovibrio* sp009773975 | 87.5 |
| CAG-485 sp002493045 | 75 |
| CAG-873 sp002490635 | 75 |
| CAG-873 sp009775225 | 75 |
| *Corynebacterium kroppenstedtii*_A | 75 |
| *Eubacterium*_J sp910586045 | 75 |
| *Gallimonas* sp910585595 | 75 |
| MGBC100798 sp910588545 | 75 |
| *Muribaculum gordoncarteri* | 75 |
| *Muribaculum* sp002473395 | 75 |

Because the absence of certain microbes may alter the disease phenotype, potentially through synergistic interactions with *C. bovis*, we identified species present in all non-protective groups but absent in A2 samples (Table 3). Among the few organisms entirely absent from the A2 group were two members of the family *Staphylococcaceae*: *Mammaliicoccus lentus* and *Staphylococcus nepalensis*.

**Table 3.** Cutaneous bacterial species detected in at least 2 samples from all 4 non-protective microbiomes and absent from the A2 microbiome.

| **Taxonomic Identification** | **Microbiome Prevalence (%)** |
| --- | --- |
| *Anaerotruncus* sp003612625 | 87.5 (B), 50 (A1+Ca, C), 37.5 (A1) |
| COE1 sp003513705 | 87.5 (B), 75 (A1), 50 (A1+Ca, C) |
| *Coproplasma* sp910578145 | 100 (B), 87.5 (C), 75 (A1), 62.5 (A1+Ca) |
| *Enterenecus* sp910588105 | 100 (A1, A1+Ca), 87.5 (B), 62.5 (C) |
| *Mammaliicoccus lentus* | 50 (A1+Ca, B), 37.5 (A1, C) |
| *Staphylococcus nepalensis* | 100 (A1), 87.5 (C), 50 (A1+Ca), 37.5 (B) |

### Fecal microbiome

Fecal sequencing yields averaged ∼15% host, ∼85% bacteria, and <1% unassigned reads. Alpha diversity did not significantly differ among microbiome groups across Observed, Shannon, or Simpson indices (Figure 6). As expected, the number of observed species in the gut was over 4-fold higher than skin samples, reflecting greater microbial richness. This was further reflected by a substantially higher proportion of bacterial versus host reads in fecal samples relative to skin.

**Figure 6.**
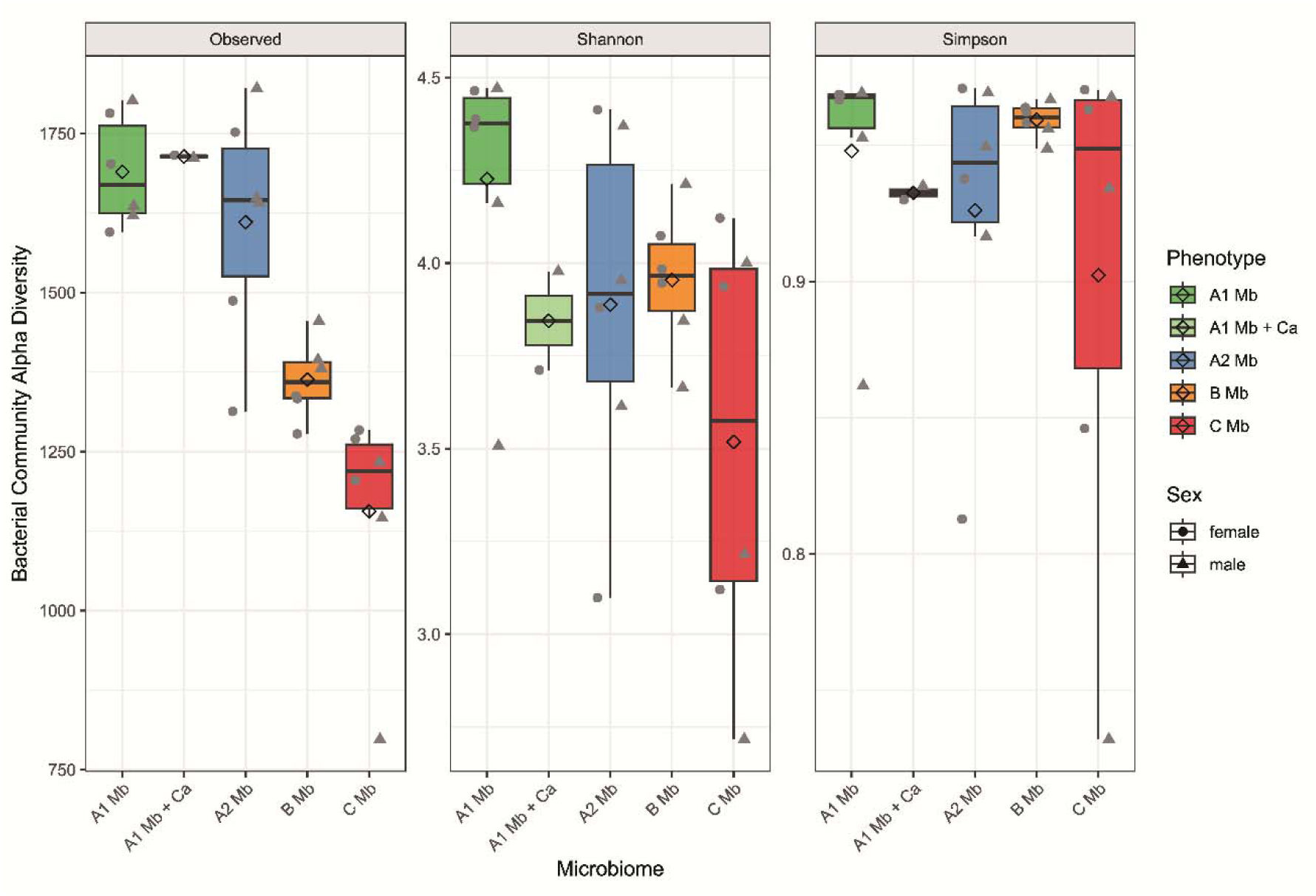
Alpha diversity of the gut microbiome across microbiome groups. Gastrointestinal bacterial alpha diversity metrics measured by Observed species richness (left facet), Shannon diversity index (middle facet), and Simpson evenness index (right facet) for pooled fecal samples across five distinct microbiome profiles: A1 Mb (dark green), A1 Mb + Ca (A1 + *Corynebacterium amycolatum*; light green), A2 Mb (blue), B Mb (orange), and C Mb (red). Individual grey data points represent individual samples, stratified by biological sex (female = circles; male = triangles), and black diamond points represent the sample mean. Box plots display the median (horizontal line), interquartile range (IQR; box limits), and minimum/maximum values (whiskers). Overall, alpha diversity of the gut microbiome did not significantly differ among microbiome groups across Observed, Shannon, or Simpson indices. There is substantially higher overall scale of observed richness in the gut (∼1,250-1,750 species) compared to cutaneous samples as seen in Figures 1 and 2.

Beta diversity metrics for the gut microbiome also demonstrated clustering among the three Vendor A groups (A1, A1+Ca, A2) consistent with the skin sample findings. However, in contrast to the skin microbiota, Vendor C samples were the most disparate when evaluated by Jaccard distance (Figure 7A), but exhibited greater community overlap with Vendor A groups when using Bray-Curtis dissimilarity (Figure 7B).

**Figure 7.**
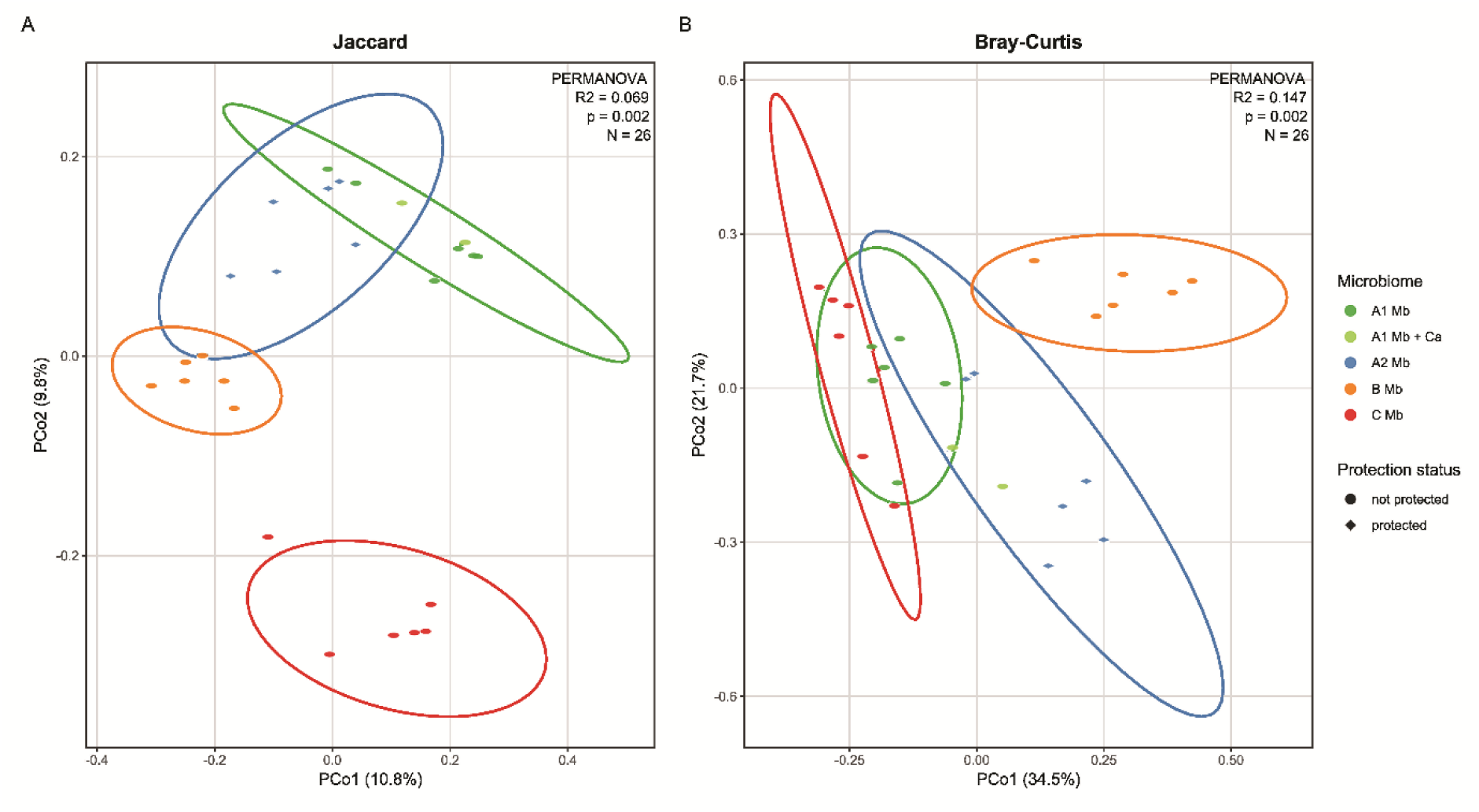
Beta diversity of the gut microbiome across microbiome groups and clinical phenotype. Principal Coordinate Analysis (PCoA) plots of pooled fecal bacterial community structures evaluated by **(A)** Jaccard distance (presence/absence) and **(B)** Bray-Curtis dissimilarity (relative abundance). Points are colored by microbiome source: A1 Mb (dark green), A1 Mb + Ca (light green), A2 Mb (blue), B Mb (orange), and C Mb (red). Symbol shapes denote disease phenotype: non-protective (circles) and protected (squares). Ellipses delineate 95% confidence intervals for each microbiome cohort, except the A1 Mb + Ca group for which only 2 samples were analyzed. PERMANOVA statistical metrics (R^2^ effect sizes and *P* values) are annotated within each panel. Note that while Vendor C gut samples form a distinct cluster away from all other groups in Jaccard distance (A), they show substantial community overlap with Vendor A groups when evaluated via Bray-Curtis dissimilarity (B).

The relative abundance of corynebacterial species was lower in the gut than on the skin. *C. amycolatum* was not detected in any of the fecal samples, despite it being a prominent member of Vendor B’s skin microbiome. Conversely, low levels of several *Corynebacterium* species were detected in the gut that were not observed on the skin, including *C. mastitidis*, *C. oculi*, and *C. resistans*. A subset of gut *Corynebacterium* remained unclassified, limiting a full comparison of cutaneous and intestinal *Corynebacterium* communities (Figure 8). Overall raw read counts for individual *Corynebacterium* species in the gut were low (Figure 9) with modest amounts of *C. bovis* present across all microbiome groups except in group A2, where raw read counts were lower than all other microbiomes.

**Figure 8.**
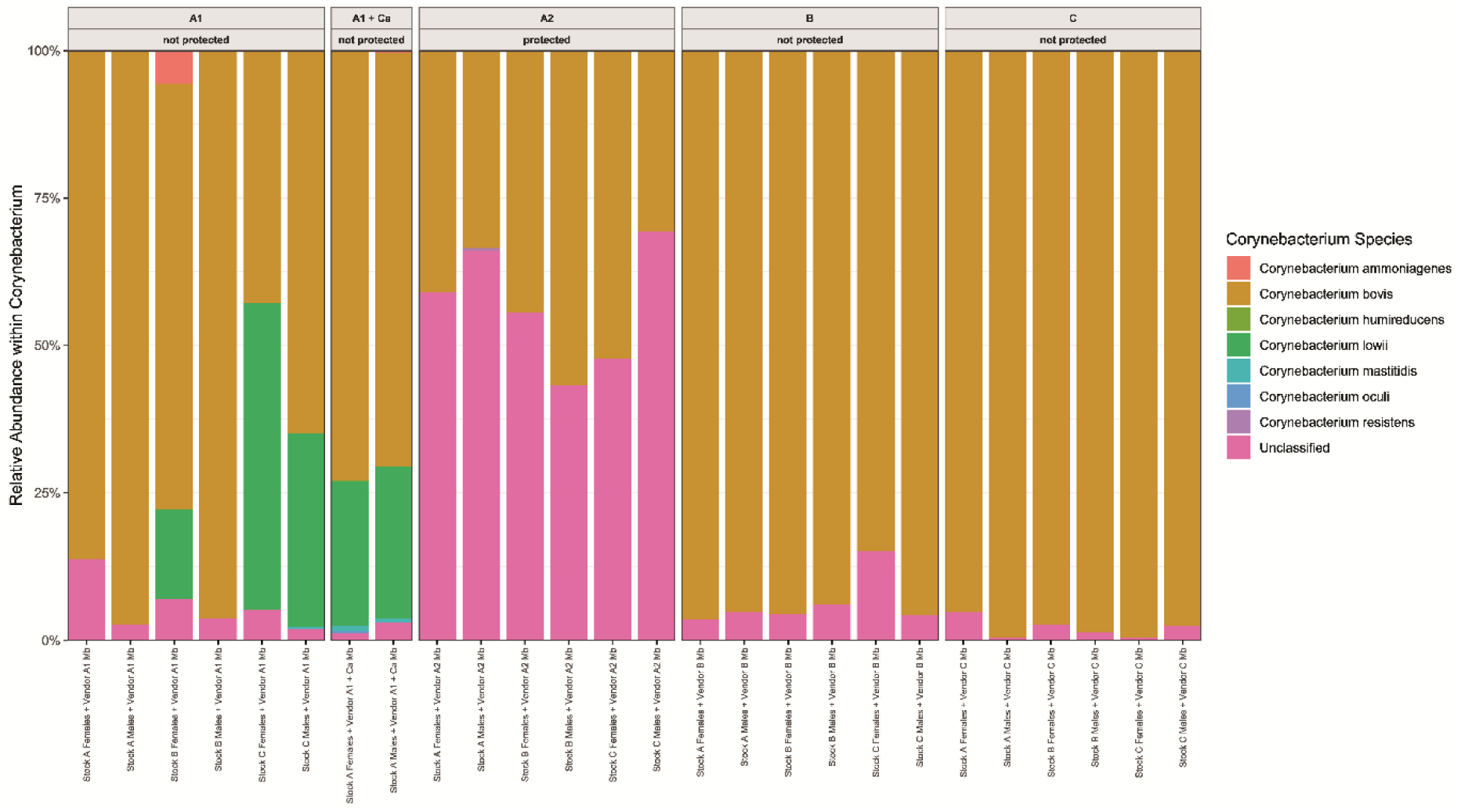
Relative abundance of *Corynebacterium* species within the gut microbiome. Taxonomic breakdown and relative abundance (%) of *Corynebacterium* species detected in fecal samples across five microbiome groups: A1, A1 + Ca (A1 + *C. amycolatum*), A2, B, and C. Clinical phenotype is annotated above each facet panel. Each vertical bar represents a single animal sample. Individual *Corynebacterium* species are indicated by fill colors, with *C. bovis* shown in gold/brown, *C. lowii* in green, and unclassified *Corynebacterium* taxa in bright pink. Note the high proportion of unclassified *Corynebacterium* reads (∼40%-70%) specifically present in the protective A2 gut flora, as well as the detection of *C. lowii* in group A1 and A1 + Ca samples.

**Figure 9.**
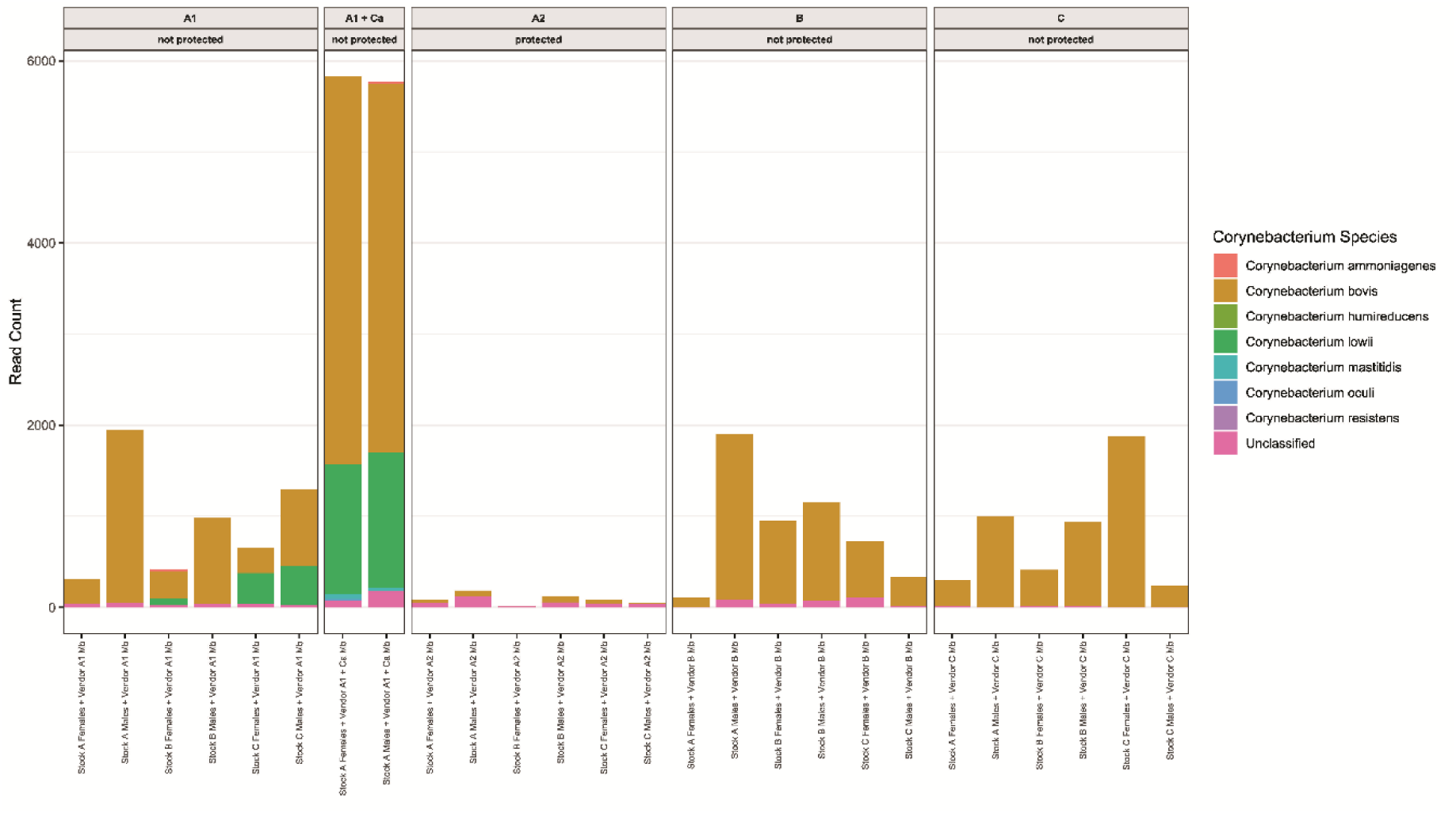
Absolute sequencing read counts for fecal *Corynebacterium* species across microbiome groups. Stacked bar plots depicting raw sequencing read counts assigned to *Corynebacterium* species in fecal samples across five microbiome groups: A1, A1 + Ca (A1 + *C. amycolatum*), A2, B, and C. Clinical phenotype status is annotated in the upper facet headers. Individual vertical bars represent individual animal samples. Species identity is indicated by fill colors, with *C. bovis* in gold/brown, *C. lowii* in green, and unclassified *Corynebacterium* taxa in bright pink. Note that while *Corynebacterium* species represented a significant proportion of the relative abundance in A2 fecal samples (Figure 8), overall absolute read counts in A2 remain extremely low (< 200 reads per sample) compared to groups A1, B, and C (∼500-2,000 reads), and A1 + Ca (∼6,000 reads).

Complete species lists were compared between the protective A2 microbiome and all other groups to identify taxa enriched or depleted in the A2 microbiome (Supplementary Tables S1 and S2). Taxa present in the nonprotective microbiomes (all groups except A2) collectively represented the depleted taxa in the protected cohort (Figure 10). Enriched species were cross-referenced with the list of unique organisms from A2 skin samples. While the One Codex reference database did not detect *C. kroppenstedtii* in the A2 gut microbiome, both *Duncaniella* species (*D. dubosii* and *D. muris*) and the skin-associated anaerobes, *B. caecimuris* and *M. gordoncarteri,* were markedly enriched in the A2 fecal samples as compared to unprotected groups. Notably, *Candidatus Arthromitus sp.* SFB-mouse-NL, also referred to as segmented filamentous bacteria (SFB), was enriched by greater than 10-fold in the A2 gut microbiome.

**Figure 10.**
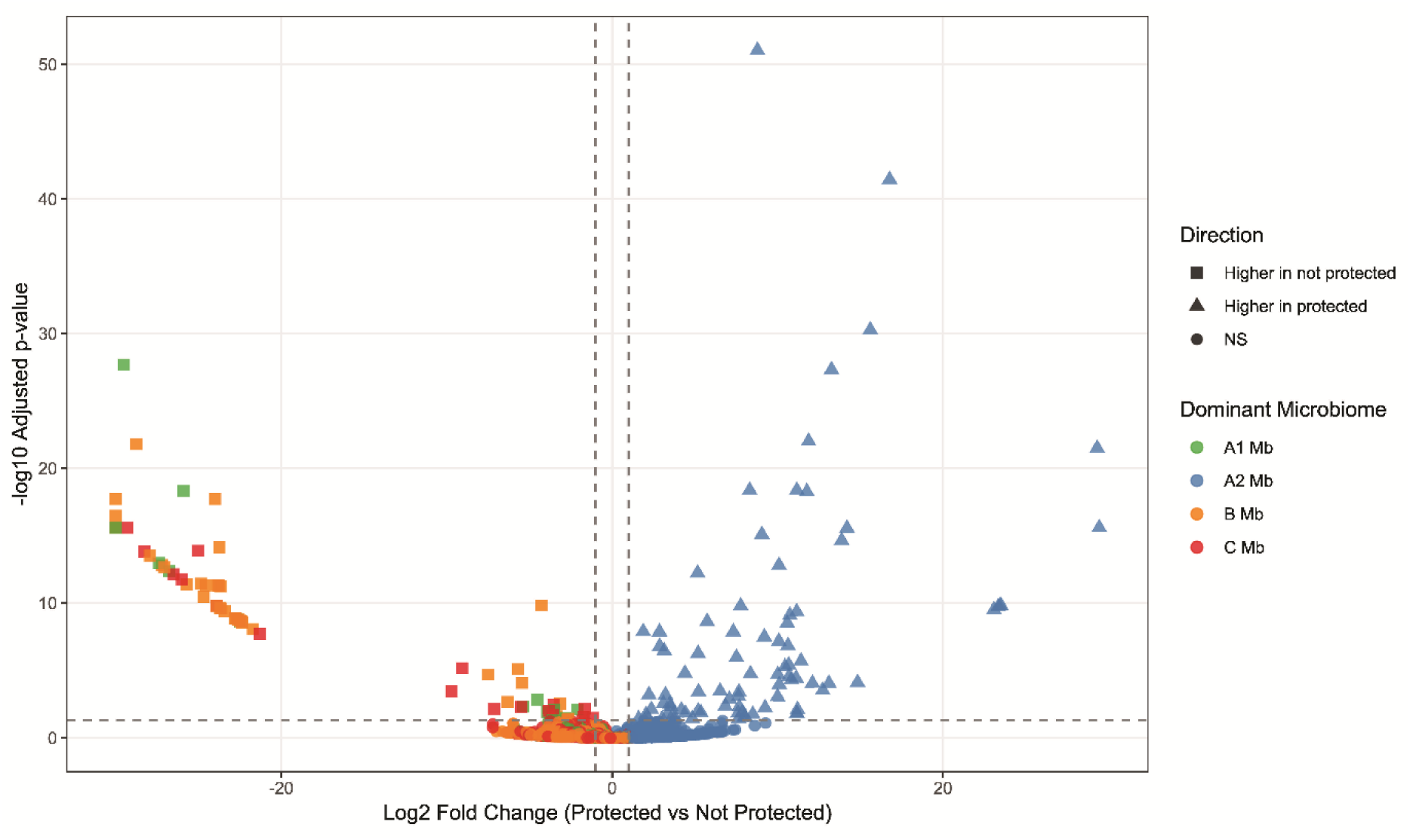
Differential abundance of gut bacterial species between protective and nonprotective cohorts. Volcano plot displaying DESeq2 differential abundance analysis of gut microbial species comparing the protective phenotype ("protected," A2 Mb) versus nonprotective phenotypes ("not protected," comprising A1 Mb, B Mb, and C Mb). The x-axis represents log_2_ fold change (positive values denote enrichment in the protective group; negative values denote enrichment in nonprotective groups), and the y-axis shows statistical significance as -log_10_ adjusted *P*-value. Dashed horizontal line indicates the threshold for statistical significance (*P*_adj_ < 0.05). Symbol shapes signify the direction of significance: triangles indicate species significantly higher in the protective group ("Higher in protected"), squares indicate species significantly higher in non-protective groups ("Higher in not protected"), and diamonds represent non-significant taxa (NS). Point colors descriptively indicate the dominant microbiome group associated with each feature: A1 Mb (green), A2 Mb (blue), B Mb (orange), and C Mb (red). Note the large cluster of significantly enriched species uniquely characterizing the protective A2 gut flora (top right blue triangles).

## Discussion

We recently demonstrated that axenic nude mice reassociated with distinct microbiomes displayed a spectrum of clinical disease and cutaneous pathology following infection with a pathogenic isolate of *C. bovis* (7894).^21^ Mice reassociated with the Vendor A2 microbiome resisted clinical disease and exhibited minimal skin pathology.^21^ The present follow-up analysis identifies several microbial features associated with this protective phenotype, including preservation of cutaneous microbial diversity after challenge, reduced relative abundance of *C. bovis*, and enrichment of candidate skin and gut taxa. Collectively, these findings suggest that protection was associated not simply with greater baseline diversity, but with the capacity of the A2 community to resist pathogen-driven disruption. The mechanism by which the A2 microbiome confers protection remains unknown and may involve several nonexclusive processes, including colonization resistance by resident taxa, absence of permissive co-colonizers, preservation of the epidermal barrier, and microbiome-mediated modulation of host innate immunity. Importantly, the presence of endogenous non-pathogenic corynebacterial species appears associated with disease resistance, with a greater diversity of corynebacterium associated with greater resistance.

*C. bovis* has previously been shown to dominate the skin microbiota of infected nude mice.^10^ We observed a similar pattern in susceptible cohorts, where increased *C. bovis* relative abundance was associated with reduced alpha diversity and more severe clinical disease. Baseline alpha diversity was comparable across most microbiome groups, whereas the protective A2 microbiome uniquely retained species richness and community evenness following *C. bovis* challenge. Thus, the distinguishing property of the A2 microbiome may be its ecological stability or resilience following pathogen exposure rather than intrinsically greater diversity before infection. This interpretation is consistent with colonization resistance, whereby an established microbial community limits pathogen expansion through occupation of ecological niches, competition for nutrients, production of inhibitory factors, or modification of the local environment. The A1 and C microbiomes had no endogenous corynebacterial species, whereas microbiomes A2 and B did. Microbiome B was associated with delayed clinical presentation in Stock A mice, microbiome A2 confers clinical protection, and the addition of a *C. amycolatum* to microbiome A1 resulted in delayed clinical onset and reduce disease severity.^21^ Colonization with non-pathogenic corynebacteria prior to *C. bovis* infection may be the primary mediator of colonization resistance, with increased corynebacterial diversity associated with greater protection.

Expansion of *C. bovis*, loss of resident organisms, and disruption of the cutaneous barrier provide a plausible ecological framework for the development of hyperkeratotic dermatitis. Dominance by a single pathogenic organism alongside reduced community diversity is characteristic of dysbiosis. While causation cannot be established from the present data, the lower relative abundance of *C. bovis* in the A2 skin microbiome aligns with previous reports of *C. bovis* expansion and concurrent depletion of skin flora in mice with active disease.^10^ It is plausible that initial *C. bovis* proliferation triggers community disruption, creating a self-amplifying cycle that enables further bacterial expansion.

Interpretation of relative abundance data requires caution. Amplicon sequencing data are compositional, such that an increase in the relative abundance of one organism may reflect its absolute expansion, a reduction in other organisms, or both.^33^ Consequently, the observed prominence of *C. bovis* cannot by itself establish an increase in absolute organism burden. Future studies incorporating quantitative culture, taxon-specific quantitative PCR, digital PCR, or spike-in standards would help distinguish absolute *C. bovis* expansion from proportional changes caused by depletion of other community members.^34^ Nevertheless, the concordance among *C. bovis* relative abundance, loss of alpha diversity, clinical disease, and histopathology supports a biologically relevant relationship between pathogen dominance and the clinical phenotype. While these data warrant cautious interpretation, the marked decrease in mean sequencing read abundance within the A2 microbiome further points to a true reduction in *C. bovis* abundance compared to other groups.

Niche competition among resident microorganisms can strongly influence pathogen colonization, particularly among phylogenetically related organisms occupying similar ecological niches.^35,36^ For example, co-colonization with resident *Corynebacterium* species in human carriers can reduce *Staphylococcus aureus* expansion by more than 70%.^37^ The greater corynebacterial diversity observed in the protective A2 skin microbiome is therefore consistent with a competitive exclusion mechanism. Resident corynebacteria could compete with *C. bovis* for essential nutrients, attachment sites, or other resources within the superficial keratin layer. Alternatively, they could modify the local physicochemical environment or inhibit *C. bovis* through direct antagonism. These mechanisms remain hypothetical and will require culture-based competition assays and controlled colonization and biochemical studies to elucidate the mechanisms involved.

Priority effects may also have contributed to protection. The sequence and timing of microbial arrival can determine subsequent community assembly and pathogen exclusion, including in germfree mouse models.^35^ The A2 microbiota was established before *C. bovis* challenge and may therefore have occupied niches that would otherwise have been accessible to the pathogen. Such an effect could explain why the community remained comparatively stable despite exposure to a high inoculation dose. Longitudinal sampling immediately before and at multiple intervals after challenge is required to determine whether protective taxa persist during early pathogen exposure and whether their presence precedes suppression of *C. bovis* expansion.

Among the candidate skin organisms, *C. kroppenstedtii* was present in all A2 animals and absent from the nonprotective cohorts. Although individual A2 samples differed somewhat in their overall *Corynebacterium* composition, the consistent detection of *C. kroppenstedtii* suggests that it may be a stable ecological feature of the protective community. It could occupy a niche similar to that required by *C. bovis*, resist displacement during challenge, produce inhibitory metabolites, or support other members of a protective consortium. However, its association with protection does not demonstrate causality. *C. kroppenstedtii* may be a direct mediator, a marker of another protective community feature, or one component of a consortium whose members are ineffective when tested individually.

*C. kroppenstedtii* is a lipophilic species with an atypical cell envelope lacking the mycolic acids characteristic of many other corynebacteria.^38^ Although poorly characterized in laboratory animal species, it is recognized as a potential opportunistic human pathogen and has been associated particularly with granulomatous mastitis and breast abscesses.^39^ Accordingly, its identification as a candidate protective organism should not be interpreted as evidence that it would be suitable for direct application as a probiotic. A safer translational objective may be to identify the ecological function, inhibitory product, or community interaction associated with its presence and determine whether that function can be reproduced by a nonpathogenic organism or a defined microbial consortium.

The results pertaining to *C. amycolatum* further support a community-level interpretation of microbiome-mediated protection. Previous work from our group identified an association between endogenous *C. amycolatum* colonization and reduced clinical expression of CAH in athymic nude mice, prompting evaluation of *C. amycolatum* as a potential protective commensal.^20^ More recently, topical administration of *C. amycolatum* before *C. bovis* challenge delayed disease onset and reduced peak clinical scores, although it did not prevent histopathologic changes or reproduce the protection observed with a nonpathogenic *C. bovis* isolate.^2^ In the present study, addition of *C. amycolatum* to the otherwise nonprotective A1 microbiome likewise did not fully reproduce the protective phenotype associated with the A2 microbiome.^21^ The relative abundance of *C. bovis* was lower in the group supplemented with *C. amycolatum*. Moreover, *C. amycolatum* was detected at only low relative abundance and was not recovered by culture despite administration of a high inoculation dose. Together, these observations argue against *C. amycolatum* as a sufficient single-organism determinant of resistance to CAH. Instead, its presence in some protective or relatively resistant colonies may reflect a broader microbial community state, an ecological interaction with other *Corynebacterium* species, or a niche that is permissive for other organisms capable of limiting *C. bovis* expansion. Thus, *C. amycolatum* remains biologically relevant as a candidate member or marker of a protective community, but its independent contribution to colonization resistance remains unresolved. Improved clinical outcomes in microbiome groups containing *C. amycolatum* further indicate that the presence of endogenous non-pathogenic corynebacterial species, rather than a single protective organism, is the primary driver of disease resistance.

Other organisms enriched in the A2 microbiome may also contribute to protection. *Muribaculum gordoncarteri* and *Duncaniella* spp., both members of the family *Muribaculaceae*, along with *Bacteroides caecimuris*, were unique to the A2 skin microbiome and enriched in A2 fecal samples. *Duncaniella muris* has been reported to reduce dextran sodium sulfate (DSS)-induced colitis and attenuate colonic shortening in axenic *Dusp6*-deficient mice.^40^ *Duncaniella dubosii* has been associated with potential T-cell immunomodulatory effects in tryptophan-supplemented mice, although a T-cell-dependent mechanism would have limited applicability in athymic nude mice.^41^ Members of the *Muribaculaceae* have been implicated in maintenance of the intestinal mucus layer, and *M. gordoncarteri* has only recently been described as a murine gut commensal.^42,43^ Similarly, *B. caecimuris* has been characterized as a keystone murine gut organism capable of influencing broader community composition.^44^ While these observations are compelling, they do not establish specific protective mechanisms on the skin of *C. bovis*-infected nude mice.

Segmented filamentous bacteria (SFB) were enriched in, although not exclusive to, the A2 gut microbiome. SFB colonization can produce strong immunostimulatory effects and is particularly associated with induction of mucosal Th17 responses.^45^ Because athymic nude mice lack normal thymus-derived T-cell populations, canonical SFB-driven adaptive responses are unlikely to fully explain protection in this model. Residual T-cell development or “leakiness” can occur in some nude mice, but the reproducibility of the protective phenotype across all three mouse stocks makes variable T-cell leakiness an unlikely sole explanation. SFB and other gut organisms may nevertheless influence epithelial signaling, antimicrobial peptide production, myeloid-cell activation, microbial metabolites, or other innate pathways. Therefore, the potential contribution of such mechanisms cannot be excluded, though gut-mediated immune protection remains hypothetical rather than established.

An alternative or complementary explanation is that susceptible microbiomes contain organisms that facilitate *C. bovis* colonization or amplify cutaneous injury. Endogenous *Staphylococcus*, *Streptococcus*, or other pathobionts could theoretically alter epidermal layers, generate inflammatory products, or otherwise create a permissive environment for *C. bovis*.^46^ While the present study did not directly establish synergism between *C. bovis* and any specific co-colonizer, *Mammaliicoccus* (formerly *Staphylococcus*) *lentus* and *Staphylococcus nepalensis* are two particularly relevant species absent from the A2 microbiome that were present in all other microbiome groups. Other *Staphylococcus* species have been reported to cause dermatitis in immunodeficient mice.^25,47^ However, the *Staphylococcaceae* family is a prominent colonizer of mouse skin, and both species also colonize other body systems. While *S. nepalensis* has been linked to exacerbating pulmonary fibrosis in mice,^48^ neither species have been implicated in causing dermatitis in immunodeficient mice. The absence of a candidate organism from sequencing data should also not be interpreted as proof of true biological absence because detection depends on organism abundance, sampling depth, DNA extraction, primer performance, and taxonomic classification. The possibility of permissive co-colonizers is therefore presented as a testable hypothesis. Pairwise and consortium-based colonization experiments are required to determine whether candidate susceptible-community organisms increase *C. bovis* burden or disease severity.

The fecal findings require separate interpretation from the cutaneous microbiome. *Corynebacterium* species commonly colonize skin and mucous membranes, and lipophilic *C. bovis* primarily inhabits the keratin layer of the skin.^46,49^ Although some corynebacteria may colonize the gastrointestinal tract, detection of cutaneous species in feces may also result from grooming and ingestion. A2-reassociated mice likely experienced less pruritus and grooming because of their reduced clinical disease, potentially decreasing ingestion of infected skin flakes and transient passage of *C. bovis* through the gastrointestinal tract. Conversely, mice in the A1+Ca group developed delayed clinical disease, with resolution occurring closer to the experimental endpoint.^21^ Increased pruritus and grooming near the time of fecal collection could therefore have contributed to the greater fecal relative abundance of *Corynebacterium* in that group. Because grooming was not quantified and fecal detection does not demonstrate gastrointestinal colonization, this explanation remains inferential. Interestingly, although skin samples from groups A1 and C lacked endogenous *Corynebacterium* species, fecal samples from these groups revealed non-*C. bovis Corynebacterium*.

Unlike the susceptible skin microbiomes, the fecal microbiomes did not show overt dysbiosis characterized by dominance of a single organism. Subtler differences in gut community structure or function, however, cannot be excluded. Moreover, endpoint fecal sampling cannot determine whether gut-community differences preceded cutaneous disease, developed in response to disease, or reflected altered grooming, food intake, inflammation, or other behavioral changes. Longitudinal fecal and skin sampling from the same individual animals would help establish the temporal relationship among gut-community composition, cutaneous *C. bovis* expansion, and onset of clinical disease.

Several limitations should be considered when interpreting these findings. Sample selection was restricted. Skin swab analysis included only 1 of the 3 mouse stocks, and fecal samples were pooled and limited to *C. bovis*-infected animals. Pooling prevented assessment of individual variability and precluded direct linking of fecal microbial features with each animal’s clinical score, skin microbiome, or pathology. The study was not statistically powered to detect differences between *C. bovis*-infected animals (n=6/group) and negative controls (n=2/group). The findings should therefore be viewed as exploratory associations and a basis for hypothesis generation rather than definitive identification of protective or permissive taxa. Cage, cohousing, and coprophagy effects may influence murine microbiome studies because animals sharing a cage are exposed to a common microbial environment and may not represent fully independent microbiological replicates.^50^ Skin and fecal samples were analyzed by different sequencing providers. Although efforts were made to harmonize the results, differences in DNA extraction, primer selection, sequencing platform, reference database, and bioinformatic pipeline can affect taxonomic profiles and diversity estimates. Direct quantitative comparison between skin and fecal datasets should therefore be avoided. The taxonomic assignments should be interpreted within the resolution of the sequencing method used. Species-level assignments, particularly for closely related corynebacteria, should ideally be confirmed by species-specific PCR, culture with isolate sequencing, targeted sequencing of additional loci, or whole-genome sequencing. Detection of microbial DNA also does not establish organism viability, stable colonization, or functional activity. Finally, despite administration of a high inoculation dose, *C. amycolatum* was not recovered by skin culture from the A1+Ca group. In contrast, *C. lowii* was detected in both the skin and gut microbiomes and was the predominant corynebacterial species after *C. bovis*. Although *C. lowii* was not identified on the skin of the A1 group, it was detected in some fecal samples from stocks B and C, for which skin swabs were not analyzed. It is possible that *C. lowii* did not establish detectable skin colonization in stock A mice under standard cohousing conditions but occupied a suitable niche following topical inoculation of *C. amycolatum*. Alternatively, inoculation may have transiently perturbed the existing community without stable establishment of *C. amycolatum*. Because culture and sequencing results were discordant, the effect of the A1+Ca intervention cannot be attributed confidently to the intended organism. However, the attenuated disease presentation in A1+Ca animals, despite minimal recovery of *C. amycolatum*, further supports the hypothesis that pre-existing colonization by corynebacterial species, rather than the presence of a single protective organism, is a primary driver of resistance to *C. bovis*.

The present study identifies candidate organisms, mechanisms, and ecological patterns but does not distinguish whether protection is mediated by an individual taxon or by the A2 community. A staged experimental approach would help establish causality. Future experiments should confirm absolute *C. bovis* burden and longitudinal community dynamics and additional studies could compare transfer of the complete A2 microbiota with supplementation using *C. kroppenstedtii*, other individual candidates, or defined A2-derived consortia. Demonstrating necessity will also be important; selective depletion of *C. kroppenstedtii* from an otherwise protective A2 community may be more informative than supplementation alone. Conversely, addition of candidate permissive organisms to the A2 community could test whether protection can be disrupted.

In summary, resistance to *C. bovis*-associated hyperkeratotic dermatitis was associated with preservation of cutaneous microbial community structure, reduced proportional dominance of *C. bovis*, and the presence of several candidate skin and gut organisms. The findings are consistent with three nonexclusive hypotheses: competitive exclusion or priority effects within the skin microbiome, absence of microbial partners that facilitate *C. bovis* pathogenesis, and modulation of host innate or epithelial responses by the skin or gut microbiota. *C. kroppenstedtii* is a prominent candidate for causal investigation, but its presence may represent a broader protective consortium rather than a single-organism effect. Longitudinal studies incorporating absolute microbial quantification, standardized sequencing, replicate cages, defined-community transfers, and targeted mechanistic analyses will be required before microbiome-based strategies for preventing *C. bovis*-associated hyperkeratosis can be developed. Greater diversity among endogenous corynebacteria may be key to conferring protection against *C. bovis*-induced clinical disease in nude mice.

## Supporting information

Supplemental Figures for Skin Microbiome

Supplemental Tables for Gut Microbiome

## Acknowledgements

We acknowledge Callista Huang, Nimisha Pattada, Abigail Michelson, Glory Leung, Juliette Wipf, Melissa Nashat, Felix Wolf, Sarah Paisner, and the staff from the Laboratory of Comparative Pathology for assistance with various technical components of the study. Generative artificial intelligence (ChatGPT, GPT-5.3, OpenAI and Gemini 3 Flash, Google) was used to assist with drafting and improving clarity of the manuscript text. All content was reviewed and finalized by the authors.

## Data access

Datasets and research notes are available at MSK.

## Nonstandard Abbreviations

CAH: *Corynebacterium*-associated hyperkeratosis
CFU: colony-forming unit
dpi: days post-inoculation
MSK: Memorial Sloan Kettering.

