## Supplemental Figures for Skin Microbiome for "Skin and Gut Microbiome Features Associated with Resistance to *Corynebacterium bovis*-Associated Disease in Nude Mice (*Mus musculus*)"

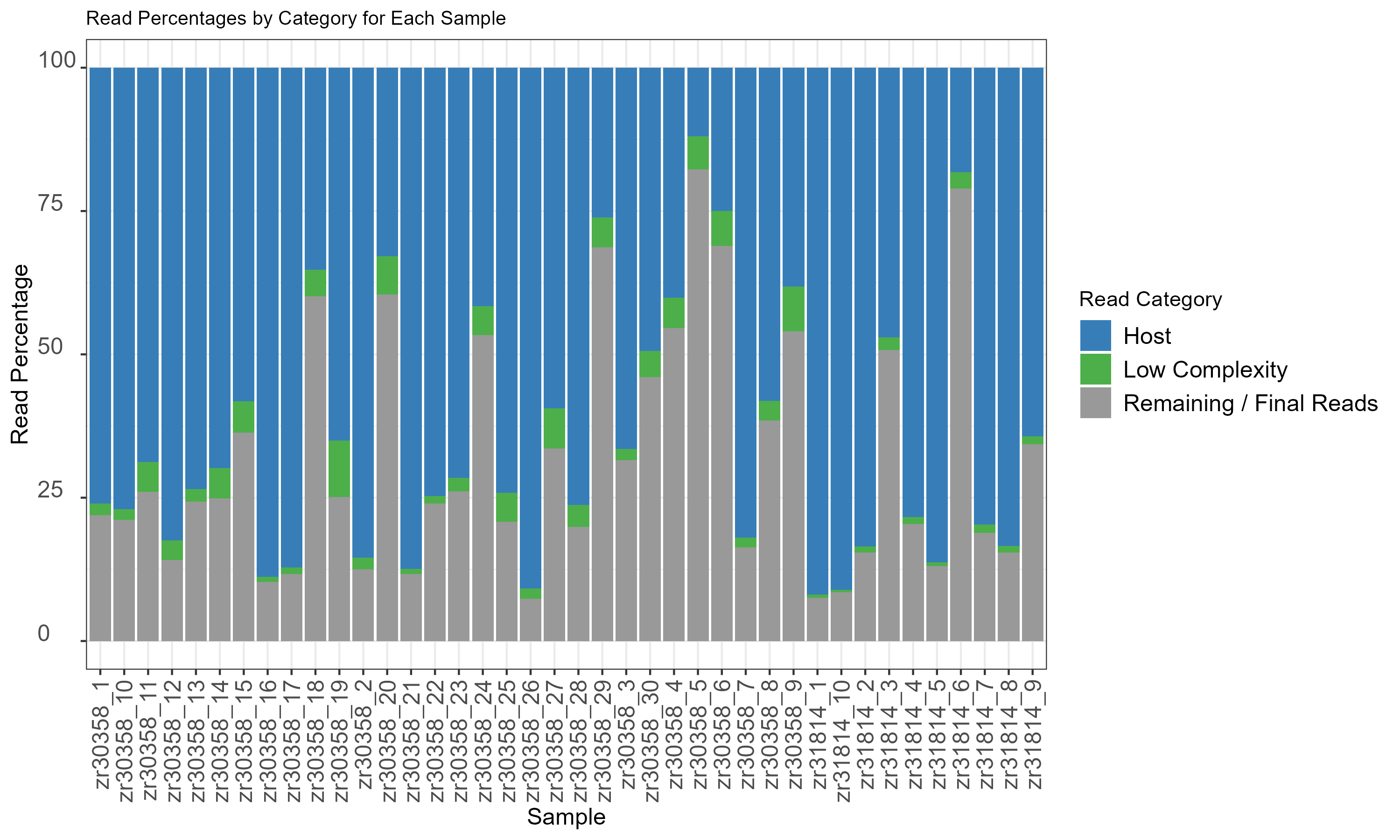

**Figure S1. Assignment of Sequenced Read Percentage in Cutaneous Samples.** Stacked bar charts showing the percentage of sequencing reads categorized as host-derived (*Mus musculus*, in blue), bacteria (in gray), or unassigned (low complexity, in green). The average host read percentage is ~29%.

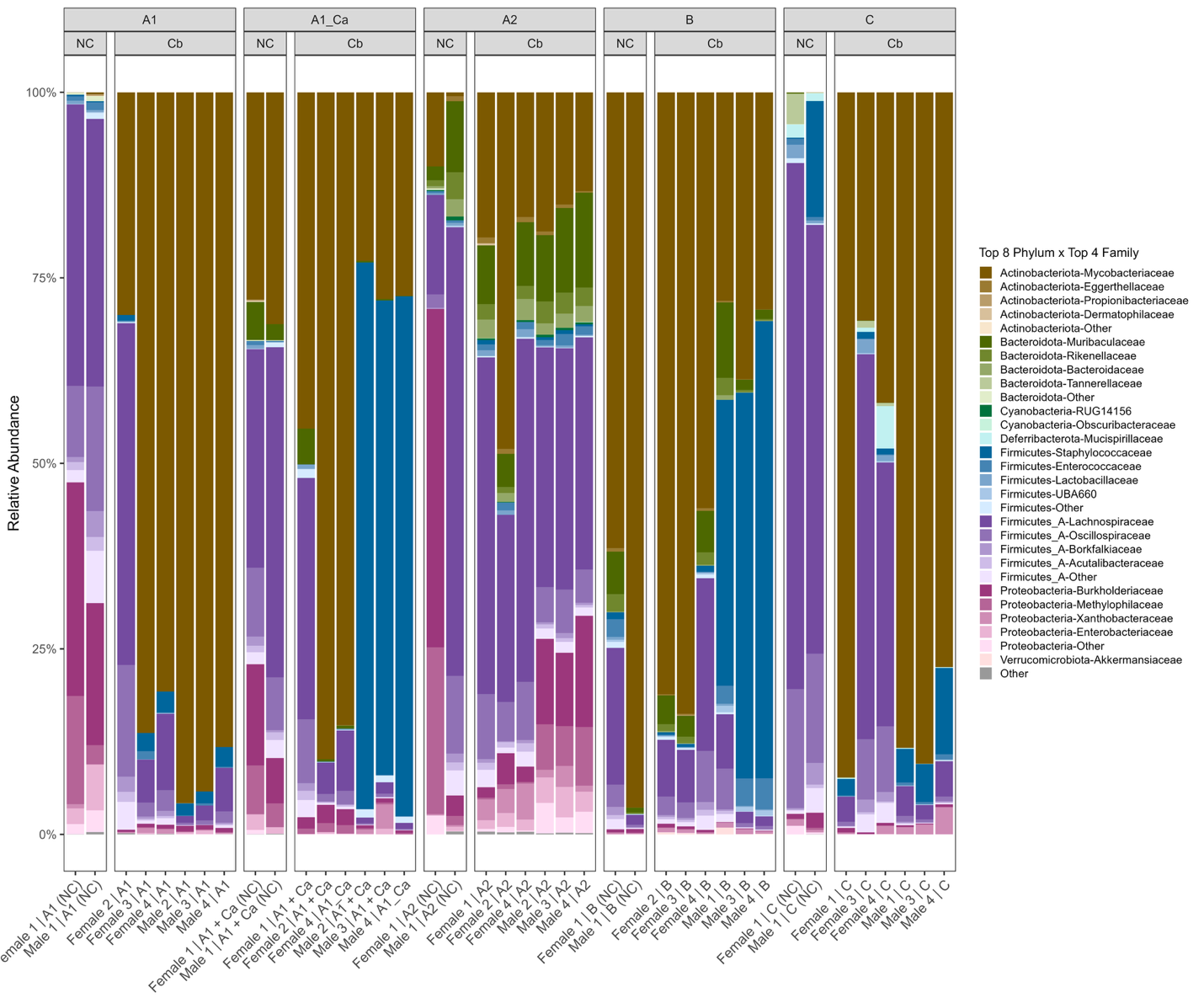

**Figure S2. Relative abundance of cutaneous bacterial phyla and family by microbiome group and infection status.** Stacked bar plots of bacterial relative abundance at phylum (top 8) and family (top 4) levels. Each bar represents one animal, grouped by microbiome source and stratified by infection status. Taxa below the display threshold are designated “Other.” NC = uninfected negative controls; Cb = *C. bovis*-infected.

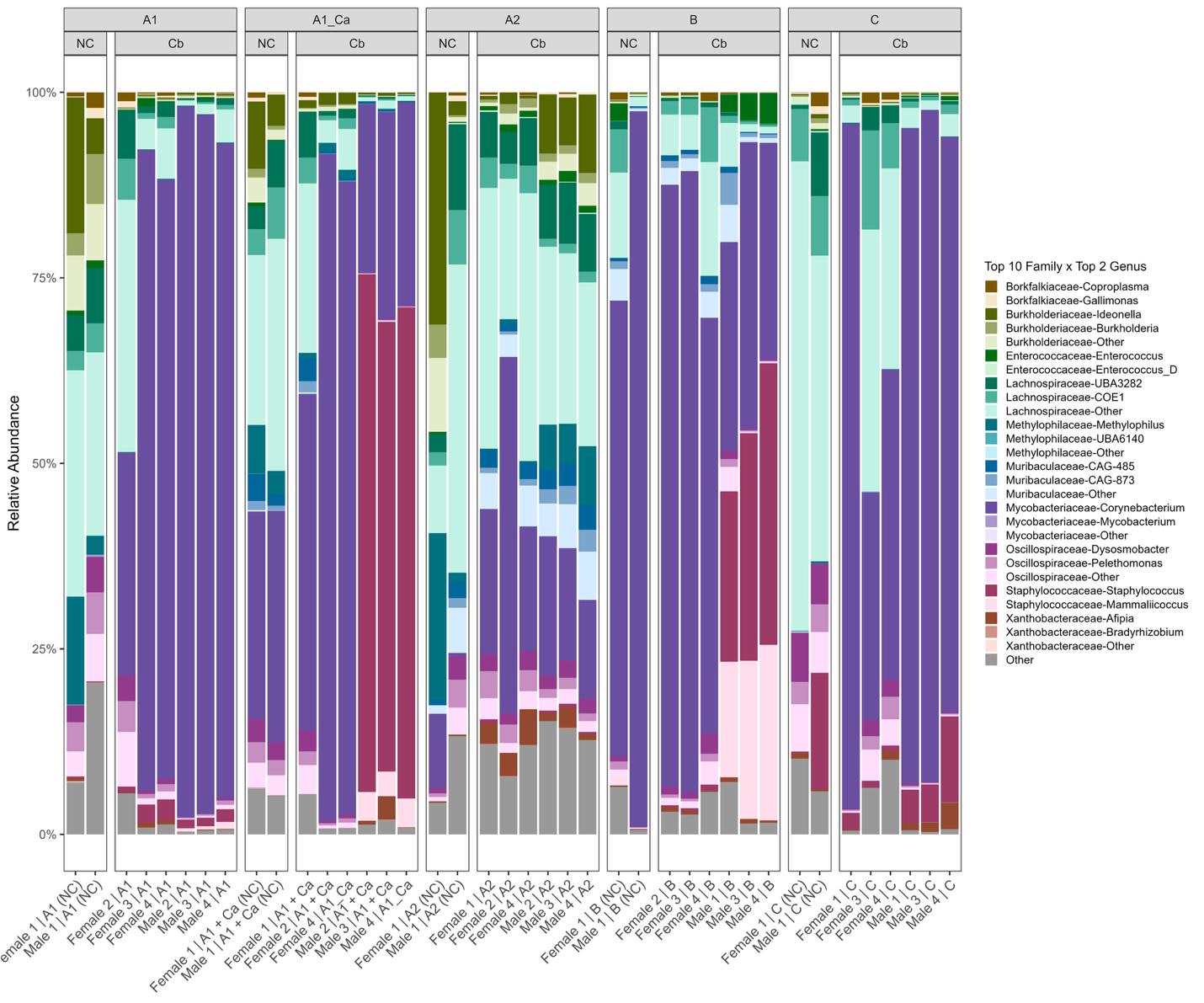

**Figure S3. Relative abundance of cutaneous bacterial family and species by microbiome group and infection status.** Stacked bar plots of bacterial relative abundance at the familial (top 10) and species (top 2) levels. Each bar represents one animal, grouped by microbiome source and stratified by infection status. Taxa below the display threshold are designated “Other.” NC = uninfected negative controls; Cb = *C.bovis*-infected.

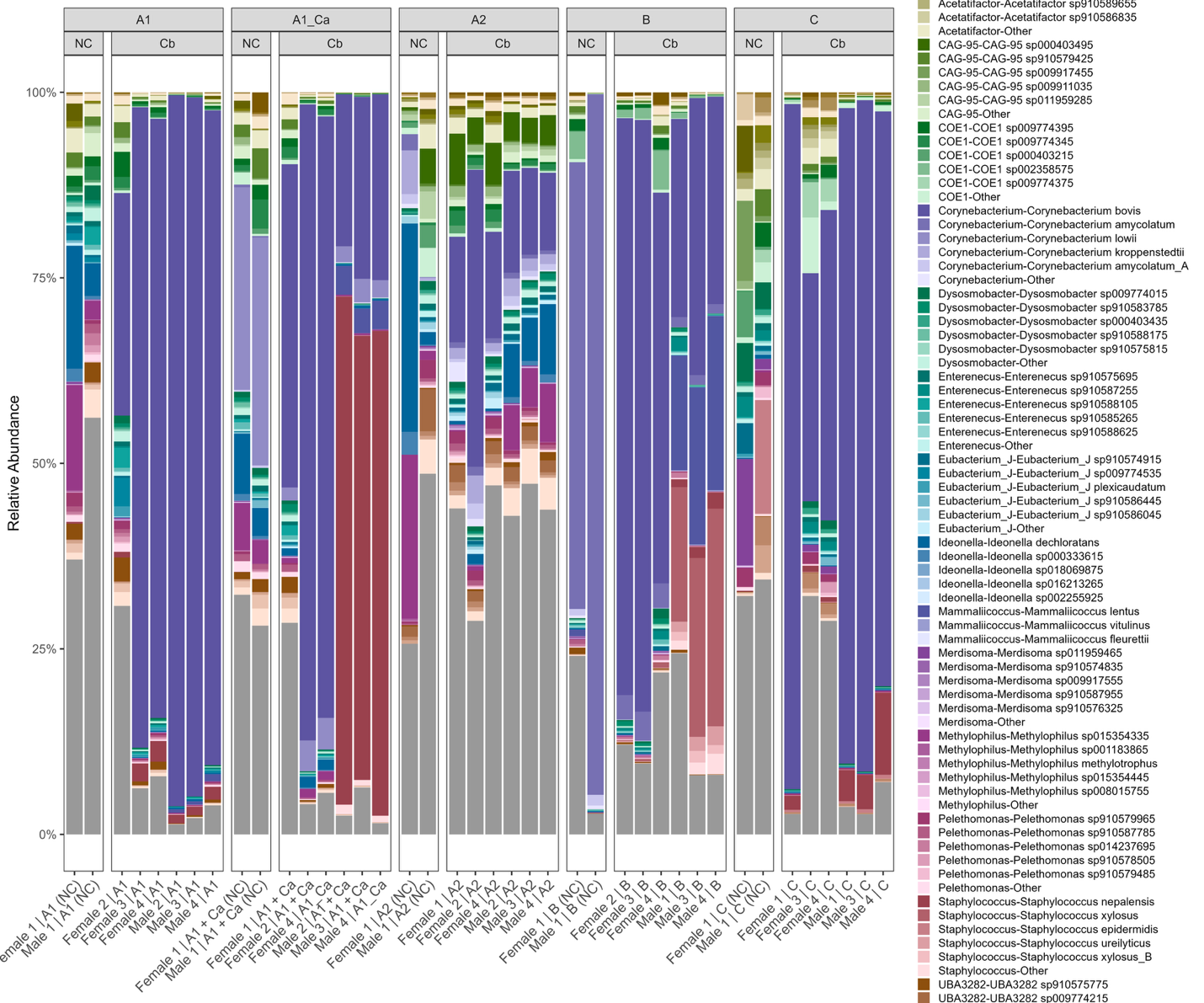

**Figure S4. Relative abundance of cutaneous bacterial species by microbiome group and infection status.** Stacked bar plots of bacterial relative abundance by genus. Each bar represents one animal, grouped by microbiome source and stratified by infection status. Taxa below the display threshold are designated “Other.” NC = uninfected negative controls; Cb = *C.bovis*-infected.
