## Supplemental Tables for Gut Microbiome for "Skin and Gut Microbiome Features Associated with Resistance to *Corynebacterium bovis*-Associated Disease in Nude Mice (*Mus musculus*)"

**Table S1.** Fecal Microbes Depleted in the Protected Microbiome after *C. bovis* challenge

| **Name** | **Rank** | **log2FoldChange** | **padj** | **baseMean** |
| --- | --- | --- | --- | --- |
| Lawsonibacter MGBC131434 | species | -6.52E-06 | 4.28E-02 | 10326.83 |
| Anaerostipes | genus | -5.10E-06 | 2.38E-02 | 88.46 |
| Corynebacterium bovis | species | -4.61E-06 | 7.09E-09 | 808.72 |
| Sporomusa | genus | -4.21E-06 | 5.46E-03 | 4.68 |
| Lawsonibacter MGBC000555 | species | -4.19E-06 | 1.38E-02 | 17856.38 |
| unclassified Anaerotruncus | no rank | -4.02E-06 | 9.64E-03 | 3064.15 |
| Borreliella | genus | -3.16E-06 | 3.47E-02 | 4.83 |
| unclassified Lachnospiraceae | no rank | -2.63E-06 | 3.20E-03 | 89263.78 |
| Caulobacter | genus | -2.32E-06 | 3.55E-02 | 5.18 |
| Dictyoglomus | genus | -2.23E-06 | 3.33E-02 | 2.38 |
| Sutterellaceae | family | -2.04E-06 | 1.00E-05 | 78.02 |
| unclassified Peptococcaceae | no rank | -1.59E-06 | 1.05E-02 | 1076.94 |
| Burkholderiales | order | -1.57E-06 | 1.36E-04 | 1816.37 |
| Enterobacter | genus | -1.49E-06 | 9.65E-03 | 261.86 |
| Ureaplasma MGBC165308 | species | -1.21E-06 | 7.37E-03 | 7.10 |
| Enterobacter cloacae complex | species group | -1.07E-06 | 3.23E-03 | 246.40 |
| Firmicutes bacterium UBA1702 | species | -1.04E-06 | 1.21E-02 | 12.94 |
| Parabacteroides | genus | -1.03E-06 | 4.97E-03 | 41263.99 |
| Staphylococcus nepalensis | species | -5.70E-07 | 6.08E-06 | 2008.10 |
| Eubacterium sp. AM47-9 | species | -4.70E-07 | 7.17E-03 | 35.74 |
| Dorea MGBC118412 | species | -4.17E-07 | 2.48E-17 | 115.48 |
| Lachnospiraceae bacterium UBA3273 | species | -2.92E-07 | 4.93E-14 | 70.53 |
| Sporofaciens musculi | species | -2.92E-07 | 5.46E-14 | 430.32 |
| Bacteroides thetaiotaomicron VPI-5482 | strain | -2.75E-07 | 5.29E-22 | 2272.79 |
| Ruminococcus MGBC117665 | species | -2.40E-07 | 5.88E-10 | 122.74 |
| Acetatifactor MGBC140221 | species | -2.37E-07 | 9.72E-23 | 3991.81 |
| Eubacterium MGBC120354 | species | -2.30E-07 | 5.43E-12 | 205.65 |
| Dorea MGBC107888 | species | -2.29E-07 | 5.27E-15 | 290.87 |
| [Clostridium] symbiosum ATCC 14940 | strain | -2.29E-07 | 1.14E-17 | 50.03 |
| Acetatifactor MGBC100080 | species | -2.15E-07 | 1.11E-08 | 22.77 |
| Mucispirillum | genus | -2.13E-07 | 5.64E-15 | 147.08 |
| Dorea sp000403475 | species | -2.01E-07 | 2.24E-10 | 57.58 |
| Mammaliicoccus lentus | species | -2.00E-07 | 7.00E-06 | 66.41 |
| Ruminococcus MGBC100158 | species | -2.00E-07 | 1.87E-08 | 26.61 |
| Staphylococcus aureus UCIM6147 | strain | -1.87E-07 | 5.05E-07 | 38.83 |
| [Clostridium] symbiosum WAL-14163 | strain | -1.82E-07 | 3.68E-10 | 17.23 |
| Lachnospiraceae bacterium UBA3401 | species | -1.78E-07 | 3.78E-16 | 22683.56 |
| Burkholderiales bacterium 1_1_47 | species | -1.77E-07 | 4.93E-14 | 59.63 |
| Kineothrix sp000403275 | species | -1.76E-07 | 5.17E-12 | 665.82 |
| Paramuribaculum MGBC114255 | species | -1.75E-07 | 2.31E-14 | 34122.96 |
| Paramuribaculum MGBC163836 | species | -1.68E-07 | 4.93E-14 | 12686.86 |
| Lachnospiraceae bacterium UBA3404 | species | -1.68E-07 | 4.93E-14 | 20751.80 |
| Bacteroides sp. AM23-12 | species | -1.67E-07 | 5.13E-14 | 1268.28 |
| Roseburia MGBC113951 | species | -1.67E-07 | 5.13E-14 | 64.54 |
| Duncaniella MGBC102972 | species | -1.67E-07 | 4.93E-14 | 66886.89 |
| Bacteroides thetaiotaomicron dnLKV9 | strain | -1.67E-07 | 3.03E-13 | 200.93 |
| Lachnospiraceae bacterium UBA7093 | species | -1.66E-07 | 1.60E-08 | 47.29 |
| Duncaniella MGBC104426 | species | -1.66E-07 | 5.99E-05 | 126.18 |
| Flavonifractor plautii | species | -1.66E-07 | 2.42E-08 | 20.53 |
| Parasutterella excrementihominis YIT 11859 | strain | -1.66E-07 | 6.46E-14 | 105.88 |
| Lachnospiraceae bacterium MD308 | species | -1.66E-07 | 5.13E-14 | 1583.69 |
| Bacteroides thetaiotaomicron CAG:40 | species | -1.59E-07 | 3.27E-12 | 658.01 |
| Acetatifactor MGBC146413 | species | -1.55E-07 | 4.93E-14 | 19298.75 |
| Eubacterium plexicaudatum ASF492 | strain | -1.06E-07 | 2.94E-15 | 2706.29 |
| Ligilactobacillus animalis KCTC 3501 = DSM 20602 | strain | -1.04E-07 | 8.18E-06 | 40.37 |
| Lachnoclostridium sp. UBA7122 | species | -8.14E-08 | 4.93E-14 | 42.84 |
| Eubacterium MGBC166763 | species | -7.84E-08 | 7.79E-16 | 8643.46 |
| Acutalibacter MGBC000522 | species | -7.51E-08 | 8.78E-14 | 309.37 |

**Table S2.** Fecal Microbes Enriched in the Protected Microbiome after *C. bovis* challenge

| **Name** | **Rank** | **log2FoldChange** | **padj** | **baseMean** |
| --- | --- | --- | --- | --- |
| Bacteroides caecimuris | species | 20.831 | 3.32E-14 | 4922.050 |
| Alistipes MGBC143807 | species | 19.531 | 4.77E-12 | 2181.115 |
| Bacteroidales bacterium M14 | species | 18.226 | 5.05E-07 | 1476.998 |
| Duncaniella freteri | species | 16.970 | 2.61E-09 | 428.857 |
| Kineothrix MGBC158835 | species | 16.492 | 6.95E-14 | 635.285 |
| Lactobacillus johnsonii N6.2 | strain | 16.186 | 6.24E-13 | 531.782 |
| Porphyromonadaceae bacterium UBA7092 | species | 15.428 | 1.56E-07 | 165.820 |
| Muribaculum gordoncarteri | species | 15.270 | 4.33E-07 | 152.871 |
| Porphyromonadaceae bacterium UBA7068 | species | 15.227 | 1.18E-09 | 157.637 |
| Porphyromonadaceae bacterium UBA7206 | species | 15.151 | 3.36E-11 | 164.113 |
| Porphyromonadaceae bacterium UBA3321 | species | 15.017 | 4.24E-05 | 185.287 |
| Porphyromonadaceae bacterium UBA3327 | species | 14.597 | 1.43E-06 | 100.502 |
| Escherichia coli | species | 13.418 | 1.62E-07 | 61.965 |
| Bacteroides acidifaciens JCM 10556 | strain | 13.380 | 4.20E-11 | 6508.972 |
| Porphyromonadaceae bacterium UBA7222 | species | 13.366 | 4.89E-03 | 70.641 |
| Porphyromonadaceae bacterium UBA3266 | species | 13.321 | 3.74E-05 | 47.626 |
| Bacteroides acidifaciens | species | 13.129 | 2.33E-12 | 14780.002 |
| Porphyromonadaceae bacterium UBA7207 | species | 13.128 | 5.86E-03 | 61.279 |
| Bacteroides sp. UBA3406 | species | 12.814 | 1.77E-03 | 729.340 |
| Candidatus Arthromitus sp. SFB-mouse-NL | species | 12.588 | 1.03E-02 | 45.717 |
| Porphyromonadaceae bacterium UBA3276 | species | 12.157 | 1.55E-04 | 23.218 |
| Porphyromonadaceae bacterium UBA7216 | species | 12.147 | 9.47E-04 | 25.610 |
| Adlercreutzia rubneri | species | 11.297 | 5.87E-04 | 13.939 |
| Lactobacillus johnsonii | species | 11.011 | 7.83E-03 | 1032.537 |
| Bacteroides ovatus | species | 10.948 | 1.02E-07 | 10.938 |
| Faecalibacterium sp. An77 | species | 10.546 | 2.60E-06 | 16.642 |
| Muribaculum intestinale | species | 10.515 | 1.32E-02 | 666.072 |
| Lepagella muris | species | 10.273 | 7.58E-08 | 1966.969 |
| Bacteroidales bacterium M13 | species | 10.246 | 6.44E-05 | 3951.236 |
| Desulfovibrio MGBC129232 | species | 10.008 | 3.61E-06 | 3205.495 |
| Porphyromonadaceae bacterium UBA3284 | species | 9.870 | 7.14E-04 | 60.111 |
| Duncaniella dubosii | species | 9.791 | 5.07E-04 | 6338.689 |
| Paramuribaculum intestinale | species | 9.770 | 1.54E-06 | 8974.066 |
| Bacteroidales bacterium M1 | species | 9.407 | 1.20E-03 | 2129.124 |
| unclassified Candidatus Arthromitus | no rank | 8.810 | 4.25E-42 | 633.606 |
| Prevotella sp. MGM2 | species | 8.600 | 2.76E-03 | 938.665 |
| Duncaniella muris | species | 8.462 | 5.24E-03 | 5541.692 |
| Prevotella MGBC143808 | species | 8.265 | 1.61E-04 | 694.596 |
| Bacteroidales bacterium M6 | species | 7.388 | 5.56E-05 | 2079.414 |
| Ligilactobacillus murinus DSM 20452 = NBRC 14221 | strain | 5.837 | 4.73E-02 | 1234.270 |
| Porphyromonadaceae bacterium UBA7149 | species | 5.807 | 1.44E-02 | 142.884 |
| Muribaculum | genus | 5.142 | 1.41E-04 | 1414.354 |
| Phocaeicola | genus | 5.018 | 4.92E-05 | 4.555 |
| Lactobacillus | genus | 4.962 | 7.21E-11 | 293.040 |
| Barnesiella | genus | 4.027 | 2.89E-03 | 36.598 |
| Duncaniella | genus | 3.965 | 9.69E-03 | 12810.429 |
| Lactobacillales | order | 3.752 | 1.36E-04 | 728.576 |
| Bacteroidaceae | family | 3.384 | 1.21E-03 | 1171.466 |
| Escherichia | genus | 3.080 | 2.13E-06 | 22.877 |
| unclassified Cyanobacteriota | no rank | 2.874 | 3.31E-06 | 10.706 |
| Capnocytophaga | genus | 2.872 | 1.93E-02 | 7.243 |
| Parvibacter caecicola | species | 2.782 | 4.30E-02 | 58.842 |
| Enterobacteriaceae | family | 2.578 | 2.98E-06 | 201.683 |
| Bacteroidales | order | 2.460 | 3.29E-02 | 184130.261 |
| Muribaculaceae | family | 2.408 | 2.98E-02 | 34857.260 |
| Prevotellaceae | family | 2.232 | 2.88E-02 | 438.153 |
| Cyanobacteriota/Melainabacteria group | clade | 1.823 | 4.69E-03 | 11.747 |
| Desulfovibrionaceae | family | 1.707 | 3.29E-06 | 35.615 |
| Dickeya | genus | 0.000 | 3.41E-02 | 3.354 |
| Bombilactobacillus | genus | 0.000 | 3.65E-03 | 1.942 |
| Cellvibrionales | order | 0.000 | 1.15E-02 | 1.727 |
| Candidatus Melainabacteria | phylum | 0.000 | 7.48E-05 | 2.133 |
| Devosiaceae | family | 0.000 | 3.35E-02 | 1.934 |
| environmental samples | no rank | 0.000 | 5.99E-03 | 4.862 |
| unclassified Oxalobacteraceae | no rank | 0.000 | 3.90E-02 | 2.532 |
| Porphyromonadaceae bacterium UBA7078 | species | 0.000 | 9.64E-03 | 8.022 |
| Lachnospiraceae bacterium UBA7143 | species | 0.000 | 1.13E-13 | 968.360 |
| Porphyromonadaceae bacterium UBA7091 | species | 0.000 | 4.88E-09 | 9.981 |
| Bacteroidales bacterium UBA714 | species | 0.000 | 1.83E-08 | 8.390 |
| Lactobacillus johnsonii NCC 533 | strain | 0.000 | 1.80E-11 | 120.281 |
| Lactobacillus johnsonii 16 | strain | 0.000 | 5.14E-09 | 8.407 |
| Pseudoflavonifractor sp. An85 | species | 0.000 | 1.89E-09 | 43.417 |
| Porphyromonadaceae bacterium UBA7180 | species | 0.000 | 9.37E-09 | 7.422 |
| Acutalibacter MGBC129708 | species | 0.000 | 1.13E-11 | 524.761 |
| Mucispirillum schaedleri ASF457 | strain | 0.000 | 8.59E-17 | 10357.086 |
| Duncaniella MGBC164402 | species | 0.000 | 4.93E-14 | 40671.076 |
| unclassified Sutterellaceae | no rank | 0.000 | 3.95E-11 | 63.399 |
